# An open-source semi-automated workflow for multi-omics sample extraction

**DOI:** 10.64898/2026.11.17.767423

**Authors:** David R. Hall, Jesse B. Trejo, Erin L. Bredeweg, Kylee L. Tate, Josie G. Eder, Priscila M. Lalli, Paul Piehowski, Tao Liu, Olga Shishkov, Kimberly DeSousa, Marina A. Gritsenko

**Affiliations:** Biological Sciences Division, Pacific Northwest National Laboratory, Richland, WA 99354, USA; Environmental Molecular Sciences Laboratory, Pacific Northwest National Laboratory, Richland, WA 99354, USA

**Keywords:** MPLEx, multi-omics, sample preparation, laboratory automation, liquid handling, Opentrons, SP3, proteomics, *Pseudomonas putida*

## Abstract

Multi-omics analysis relies on extracting metabolites, lipids and proteins from a single sample. A widely used method for this is Metabolomics, Proteomics, and Lipidomics Extraction (MPLEx), a triphasic chloroform-methanol-water extraction, but it is manually intensive and depends on skilled operators working around a fragile protein layer. Here, we present OpenMPLEx, a semi-automated MPLEx workflow for low-cost, open-source Opentrons liquid handlers. A minor change in chemistry pellets the protein, so that a robot can collect the aqueous and organic phases from above. We compared the automated workflow with two manual MPLEx protocols by liquid chromatography–mass spectrometry (LC-MS)-based proteomics, lipidomics and metabolomics. The methods were roughly equivalent and measured similar differences between growth media. We then scaled the workflow to 96-well plates and tested it on an 80-sample *Pseudomonas putida* experiment. One operator took all 80 samples from cell pellets to LC-MS data in a working week, with about 3 h of hands-on time rather than days at the bench, and quantified about 3,900 protein groups. All scripts, labware definitions and print files are open source, so that other laboratories can adapt the workflow.

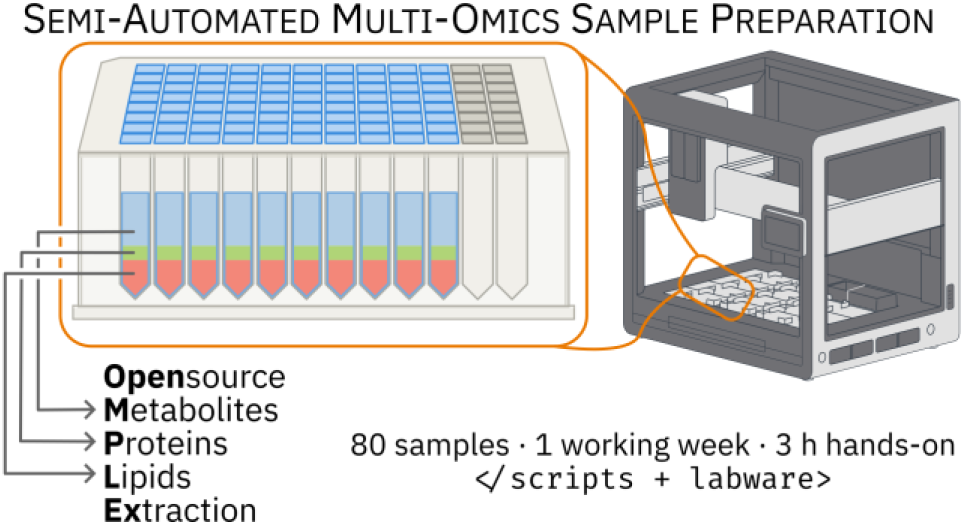

## Introduction

Integrative multi-omics analysis, capturing proteins, lipids, and polar metabolites from the same biological sample, has become central to understanding cellular systems^1^ and diseases^2^. Advances in LC-MS acquisition speed and sensitivity have made it feasible to measure thousands of analytes across these molecular classes,^3^ while modern experimental designs and statistical frameworks increasingly demand larger sample sizes.^4^ This creates a fundamental mismatch: analytical capacity has scaled, but sample preparation remains largely manual.

The Metabolomics, Proteomics, and Lipidomics Extraction (MPLEx) protocol uses a chloroform-methanol-water phase separation to partition a single sample into three fractions: an aqueous phase containing polar metabolites, an organic phase enriched in lipids, and precipitated proteins.^5^ This protocol is robust and broadly applicable to many sample types including, but not limited to, bacteria,^6^ fungi,^7^ plants,^8^ soil,^9^ and biofluids.^10^ Furthermore, MPLEx has been shown to inactivate pathogens during extraction,^11^ which makes it attractive for research on potentially higher-risk samples. However, the tri-phasic extraction is labor-intensive and requires skilled technicians to navigate the gelatinous protein interface that forms between the aqueous and organic layers.^12^

Automation of sample preparation using chloroform-methanol chemistry has been demonstrated for proteomics, where Chen et al. (2019) showed that the solvent mixture simultaneously lyses cells and precipitates proteins, and that a modified extraction order enables reliable plate-based processing on liquid handling robots.^13^ Their implementation on a fully integrated platform achieved high reproducibility for proteomic workflows. However, MPLEx and similar extractions are used across a wide range of experimental contexts, from large cohort studies to one-off pilot experiments, each with different downstream analytical requirements. A tightly integrated system optimized for a single end-to-end workflow does not readily accommodate this diversity.

Here, we present OpenMPLEx, a semi-automated adaptation of the MPLEx protocol for open-source Opentrons liquid handling robots.^14^ By design, the workflow retains off-deck centrifugation and manual plate transfers, avoiding the cost and complexity of integrated centrifuges while preserving flexibility. Protocols are organized as modular scripts, with individual functions for solvent addition, phase collection, and on-deck protein quantification by BCA, that users can adapt, recombine, or extend for their own downstream analyses. We validated the workflow against manual MPLEx using *Pseudomonas putida* grown in two media conditions, confirming comparable performance across proteomics, lipidomics, and metabolomics by LC-MS. We then scaled the workflow to 96-well plates on the Opentrons Flex and ran a designed 80-sample carbon × nitrogen experiment. One operator took all 80 samples from cell pellets to LC-MS data in a working week with approximately 3 h of hands-on time, and the data recovered the expected differences between conditions. We also describe a low-cost ventilation solution that met our institutional safety requirements for handling volatile solvents on a liquid handler. All protocols are provided as open-source Python scripts in the Supporting Information.

## Methods

### Cell culture and harvesting

For the OT-2 extraction comparison, *P. putida* KT2440 was cultured separately in LB Lennox Broth (Teknova, L9310) and Reasoner’s 2A Broth (R2A; Neogen, NCM0188A). For each medium, 20 mL in a 125 mL flask was inoculated at a 1:200 dilution and grown for 5–6 h at 28 °C with shaking at 200 rpm to an *OD*_600_ of about 1. Cultures were harvested at *OD*_600_ = 1.25 (LB) and 1.03 (R2A). For each replicate, 1.2 mL of culture was pelleted by centrifugation, washed once with 1× PBS, flash-frozen in liquid nitrogen and stored at −80 °C until processing.

For the Flex scale-up plates, *P. putida* KT2440 was grown as a single culture. A single colony was grown overnight in LB and transferred into 30 mL of modified M9 medium (6.78 g/L Na_2_HPO_4_, 3 g/L KH_2_PO_4_, 0.5 g/L NaCl, 2 mM MgSO_4_, 100 µM CaCl_2_, 18 µM FeSO_4_, 30 mM MOPS, 1 g/L Teknova vitamin mix, 20 g/L glucose and 9.24 g/L (NH_4_)_2_SO_4_), prepared by sterile filtration. The flask was grown overnight at 28 °C and 200 rpm to an *OD*_600_ of 7.15 to provide enough biomass. Cells were dispensed into 80 wells of each plate, washed in the plate with 1× PBS, pelleted by centrifugation and stored at −80 °C until processing.

For the carbon × nitrogen experiment, *P. putida* KT2440 was grown from a glycerol stock overnight in 3 mL LB, passaged into 30 mL of the modified M9 medium used for the Flex scale-up plates for acclimation, and inoculated 1:100 into three independent 125 mL flasks per condition, each containing 50 mL of medium. Cultures were incubated at 28 °C with shaking at 200 rpm. The flasks contained the same M9 base without glucose or (NH_4_)_2_SO_4_ (6.78 g/L Na_2_HPO_4_, 3 g/L KH_2_PO_4_, 0.5 g/L NaCl, 2 mM MgSO_4_, 100 µM CaCl_2_, 18 µM FeSO_4_, 30 mM MOPS and 1 g/L Teknova vitamin mix), with 30 mM glucose or *p*-coumarate as the carbon source and 18 mM NH_4_Cl or NaNO_3_ as the nitrogen source (Figure 4 A). Cells were harvested at *OD*_600_ 0.79–1.06 (Figure S1), washed twice with 1× PBS and stored at −80 °C.

### MPLEx extraction comparison

Three MPLEx workflows were compared on *P. putida* KT2440 pellets grown in LB and R2A: an automated method on an Opentrons OT-2 with on-deck pipette mixing (OT-2 Robot, described below), a manual sequential MPLEx protocol matching the OT-2 chemistry (Human Sequential), and a classic manual MPLEx protocol in which the organic phase is pipetted through the protein layer (Canonical; Figure 1). The three protocols use the same chemistry until the aqueous (metabolomics) phase is removed, so our comparisons focus on the protein and lipid fractions, where the chemistry diverges. Each method was performed in quadruplicate per medium (n = 4 × 3 methods × 2 media = 24 extractions), with two solvent-only extraction blanks per workflow (OT-2 Robot and Human Sequential; n = 4 blanks total). Briefly, thawed pellets were spiked with 2 µL UltimateSPLASH ONE (Avanti Research, REF330820) internal standard, lysed with 600 µL 2:1 CHCl₃:MeOH and 200 µL water, incubated for 10 min on ice (on the 4 °C block for the OT-2) and centrifuged at 9,000 × g for 10 min at 4 °C, and the aqueous phase was collected for metabolomics. In the sequential and OT-2 protocols, 800 µL methanol (twice the chloroform volume) was then added to the remaining protein and organic phases. The added methanol mixes with and dilutes the chloroform, which lowers the density of the organic phase so that a second centrifugation at the same speed settles the protein at the bottom of the tube. The organic phase was then collected from above for lipidomics, leaving the protein pellet. In the canonical protocol, the organic phase was collected through the protein layer, and the protein was recovered without a methanol wash. Both manual workflows were performed by a single experienced operator to minimize inter-operator variability and to provide the most demanding benchmark for the automated method. All downstream metabolomics, lipidomics, and proteomics sample preparation was processed in parallel through a common workup to ensure that any differences in protein recovery reflected the extraction step alone.

**Figure 1:**
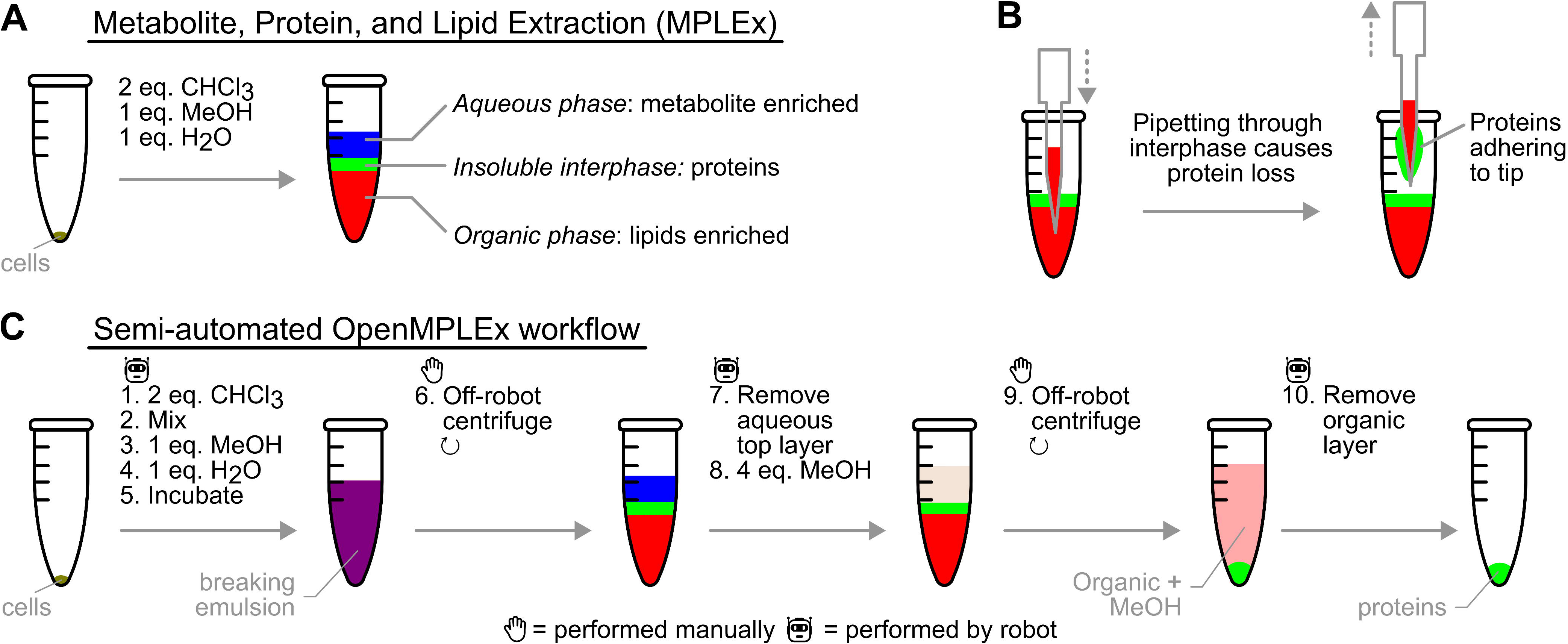
The OpenMPLEx semi-automated workflow. (A) Canonical MPLEx phase separation. (B) Pipetting the organic phase through the gelatinous protein interphase risks losing part, or all, of the protein layer to the pipette tip, a failure mode that requires visual monitoring to catch and is therefore difficult to automate. (C) OpenMPLEx instead dilutes the organic phase with a second methanol aliquot, and re-centrifuges so proteins pellet at the tube bottom, allowing clean organic-layer removal.

Protein pellets were resuspended in 8 M urea in 100 mM ammonium bicarbonate (pH 8.5) by vortexing and brief sonication, and protein concentrations were determined by Pierce BCA assay. Aliquots were reduced with 5 mM DTT (30 min, 60 °C, 500 rpm) and alkylated with 40 mM iodoacetamide (45 min, 25 °C, 500 rpm, in the dark). Samples were diluted 8-fold with 100 mM ammonium bicarbonate, and trypsin (Promega) was added at a 1:50 enzyme:protein ratio for digestion (3 h, 37 °C, 800 rpm). Digests were acidified to 1% formic acid, desalted by C18 SPE, concentrated to ∼100 µL, and quantified by BCA. Final peptide samples were diluted to 0.0125 µg/µL and loaded onto Evotip Pure (EV2001) tips using an OT-2 robot, as described below for the carbon × nitrogen experiment.^15^

Lipidomics and metabolomics sample preparation and analysis are described in the supplementary information.

### Automated MPLEx on the OT-2

The automated extraction was implemented as a custom Opentrons OT-2 protocol (Python, opentrons API v2.20) using a single-channel P1000 Gen2 pipette, a Gen2 temperature module holding samples in a 24-tube aluminum block at 4 °C, and a 3D-printed three-bottle solvent reservoir. To safely handle the chloroform and methanol used during extraction, the OT-2 was operated inside a chemical fume hood for the duration of the run. The protocol executes the chemistry described above with two off-deck pauses for incubation and centrifugation. Briefly, the robot dispenses 600 µL of 2:1 CHCl₃:MeOH into each tube using a fresh tip per sample with vigorous on-deck mixing (10 cycles of 200 µL near the tube bottom) to disrupt cell pellets, followed by addition of water to a 200 µL aqueous total volume with stepped mixing in the aqueous layer (five z-positions, two cycles each). After the first centrifugation, the aqueous (top) layer is removed by stepped aspiration that tracks the liquid surface as it descends and stops at a fixed safety margin above the empirically determined aqueous–organic interface. The robot then dispenses 800 µL of methanol to dilute the chloroform; after the second centrifugation, the organic layer is similarly recovered by stepped aspiration above the protein pellet and dispensed into glass autosampler vials. Pipette speeds for each step are listed in the scripts. Solvent bottle levels were tracked dynamically using an empirically derived height-from-volume calibration. Interface heights were determined from preliminary extractions in matched sample tubes, using the partitioning dyes methylene blue (aqueous) and oil red O (organic) to mark the layers and the depth gauge of a pair of calipers to measure them from the top of the tube (Figure S2). The dyes made automation quicker to test during development, and they make the phase boundaries easy to see when adapting the heights to other tubes or plates. The scripts, labware definitions, and STEP files for the 3D-printed labware are provided in the Supporting Information.

### Flex scale-up plates

Two 80-well plates of *P. putida* from a single culture were extracted on different days with S1 (Table 1). Protein pellets were resolubilized in 6 M guanidine-HCl, and total protein was read on deck by BCA (S2). As references, albumin was carried through S1, and the same culture was lysed directly in SDS without extraction.

**Table 1:**
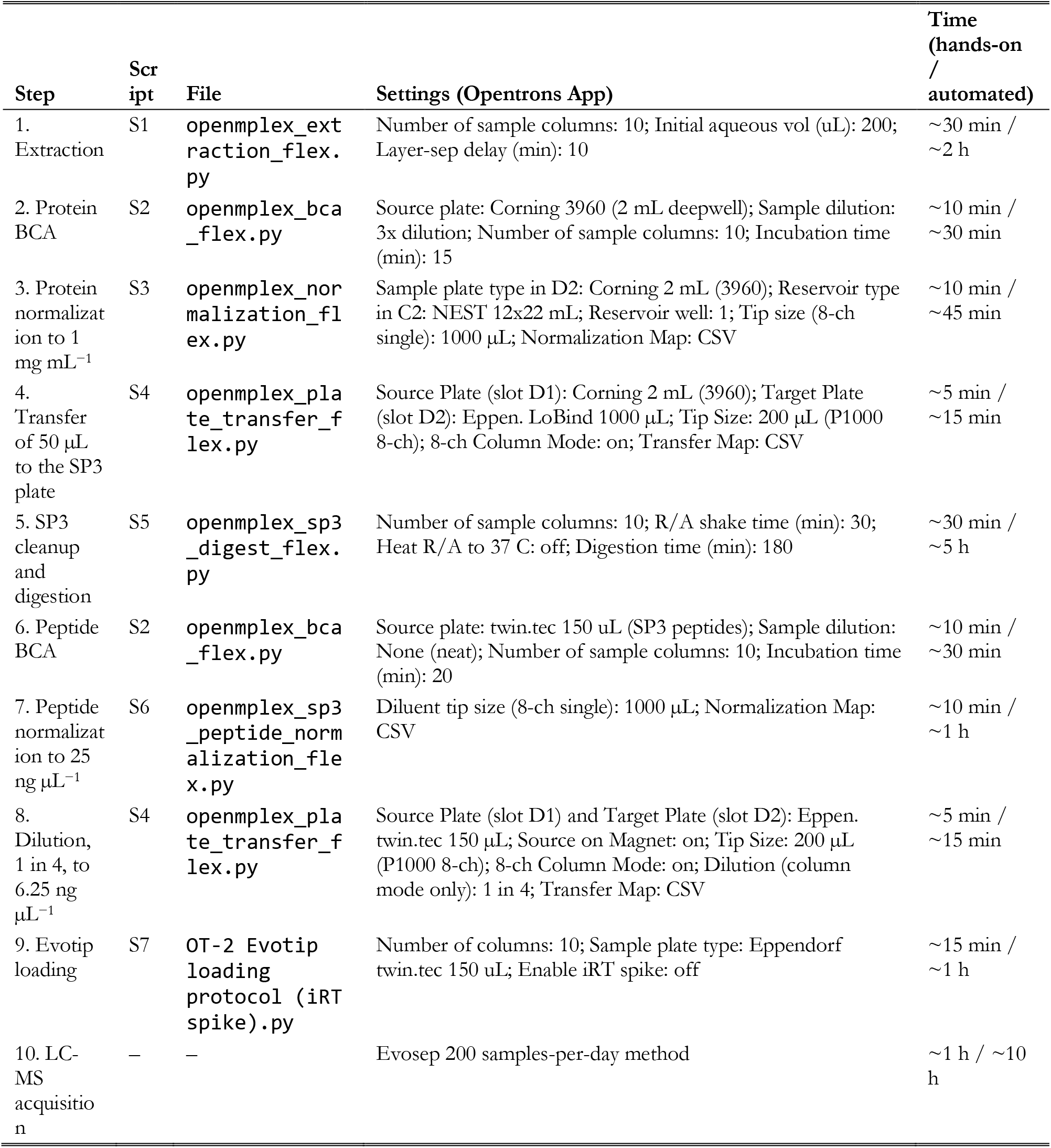
Scripts and settings for each step of the carbon × nitrogen workflow. S1–S6 ran on the Opentrons Flex and S7 on the OT-2. Settings are listed as they appear in the Opentrons App; settings not listed were left at their defaults. Map files are CSV files of per-well volumes; the preparation workbook (SI) generates them. Times are approximate, for 80 samples, and assume stock solutions are already prepared; the two dry-downs and tidying up add about 1 h of hands-on time.

### Carbon × nitrogen experiment

Frozen pellets were resuspended on ice in ice-cold LC-MS water at 39 µL per mg wet mass (25 mg mL^−1^), and 200 µL (5 mg wet cells) was loaded per well into a 96-well 2 mL plate (Corning 3960), 20 wells per condition in a balanced, randomized layout (Figure 4 B). The plate format and well assignments were kept constant from loading through Evotip loading. With the supplied scripts, wells could instead be reshuffled at the transfer to the SP3 plate (S4, in single-channel mode) or at peptide normalization (S6), both of which take a per-well map, and with changes to the scripts they could be reshuffled at other steps.

Sample preparation was automated on an Opentrons Flex (robot software 9.0), and Evotips were loaded on an Opentrons OT-2. Each step ran one of six Flex scripts (S1–S6) or the OT-2 Evotip script (S7); Table 1 lists the script and settings for each step. A preparation workbook (SI) carries the sample names from the plate map through every step, calculates the normalization volumes from the BCA readouts, and generates the input file for each script. It is filled in with this experiment’s plate map and BCA readouts as example data. The scripts and any custom labware definitions are also provided in the SI.

Samples were extracted by OpenMPLEx (S1) with ice-cold solvents, holding the plate and solvent reservoir at 4 °C. Each well received 400 µL chloroform and was shaken to lyse the cells, then 200 µL methanol and a second shake, with the 200 µL cell suspension providing the aqueous phase. After 10 min at 4 °C, the plate was centrifuged (3,000 × g, 15 min, 4 °C) and the aqueous phase was removed. A further 800 µL methanol, a second 10 min hold and a second centrifugation at the same speed pelleted the protein, and the organic phase was removed. Only the protein fraction was analyzed here; protein pellets were dried in a speed vac and stored at −80 °C.

Pellets were resolubilized in 200 µL 8 M urea, 100 mM Tris-HCl pH 8.0 (vortex, 5 min bath sonication, 30 min shaking at 15 °C), and protein was measured by BCA (Pierce, Thermo Fisher Scientific; S2) against matrix-matched BSA standards. Each well was then diluted in place with urea buffer to 1 mg mL^−1^ (S3); wells already below this concentration were not diluted, and 50 µL of every well was transferred to a 96-well Protein LoBind plate (Eppendorf; S4).

Proteins were cleaned up and digested by SP3 (S5),^16^ with full parameters in the script. The protocol was checked separately by SDS-PAGE on mouse macrophage lysate left over from another project, to confirm that proteins were captured on the beads and that peptides were released (Figure S3). Briefly, proteins were simultaneously reduced and alkylated with 10 mM TCEP and 40 mM chloroacetamide (30 min, room temperature),^17^ mixed with 500 µg Sera-Mag SpeedBeads (1:1 hydrophilic:hydrophobic; Cytiva), and aggregated onto the beads by bringing the mixture to 58% (v/v) ethanol. The beads were captured on a magnet, rinsed three times with 180 µL 80% ethanol, and digested in 150 µL 100 mM TEAB containing 1.5 µg trypsin (Promega; 1:33 enzyme:protein) for 3 h at 37 °C. The peptide supernatant (125 µL) was then drawn off the magnet-held beads into a skirted twin.tec PCR Plate 96 LoBind (Eppendorf, 0030129512). S5 as supplied includes a slower final draw adopted after this run, in which beads carried over and were removed by a second transfer off a magnet.

Peptides were dried in a speed vac, reconstituted in 60 µL 0.1% formic acid and measured by BCA (S2). They were normalized in two steps, each drawing from the source plate on a magnet: wells were first diluted to 25 ng µL^−1^ in 100 µL 0.1% formic acid on a fresh plate (S6), and 15 µL of each was then diluted 1 in 4 with 0.1% formic acid to 6.25 ng µL^−1^ (S4). For each sample, 20 µL (125 ng) was loaded onto an Evotip Pure (S7).^15^

### LC-MS/MS analysis

Peptides from the OT-2 extraction comparison (250 ng on-column) were separated on an Evosep One (Evosep) using the 60 samples-per-day method (21 min gradient) with an EV1109 performance column.^18^ Mobile phases were 0.1% formic acid in water (A) and 0.1% formic acid in acetonitrile (B; LC-MS grade solvents, Thermo Fisher Scientific). The LC was coupled to an Orbitrap Astral Zoom mass spectrometer (Thermo Fisher Scientific) operated in positive ion mode (NSI, +2.3 kV; ion transfer tube 300 °C). MS1 scans were acquired in the Orbitrap (m/z 380–980, 240,000 resolution, 500% normalized AGC, 5 ms maximum injection time) with EASY-IC lock-mass correction. DIA MS2 scans were acquired in the Astral analyzer using 200 sequential 3-Th isolation windows spanning m/z 380–980, with HCD at 25% normalized collision energy, product-ion range m/z 150–2,000, 500% normalized AGC, 7 ms maximum injection time, and a 0.6 s loop time. Carbon × nitrogen samples (125 ng on-column) were analyzed with the 200 samples-per-day method (5.6 min gradient) on an EV1107 column, with MS1 and DIA over m/z 400–800 using 4-Th isolation windows and a 3 ms maximum injection time. These samples were acquired in a randomized block order, with each of the 20 blocks containing one sample from each condition.

### Data Analysis

DIA raw files were processed with DIA-NN^19^ v2.3.2 in library-based mode using a predicted spectral library generated in silico from the *Pseudomonas putida* KT2440 reference proteome (UniProt UP000000556; downloaded 2026-02-23) supplemented with the cRAP contaminants database (camprotR, v240512)^20^. Tryptic digestion was specified with up to 1 missed cleavage and N-terminal methionine excision; carbamidomethylation of cysteine was set as a fixed modification. Peptides of 7–30 residues with precursor m/z 300–1,800 and charge states 1–4 were considered, and fragment ions were searched over m/z 200– 1,800. Match-between-runs (MBR) and RT profiling were enabled, and mass accuracy was optimized automatically. Precursor and protein group identifications were filtered to 1% FDR. Peptides mapping to cRAP entries were excluded from normalization and from quantification of protein groups not containing a contaminant. For the carbon × nitrogen experiment, precursor m/z was 400–800 and fragment m/z 150– 2,000, and mass accuracy was fixed at 10 ppm (MS2) and 4 ppm (MS1), the values recommended for the Orbitrap Astral in the DIA-NN documentation.^21^

Protein-level analysis was done in R with limma^22^ and vegan.^23^ In both experiments, MaxLFQ protein group intensities were log₂-transformed and median-centered per sample, protein groups quantified in at least 75% of the samples of at least one group were kept, per-protein fold changes were estimated with limma from one mean per group, and PCA used only protein groups quantified in every sample, without imputation. For the OT-2 extraction comparison, the same matrix was used to partition variance among media, extraction method and their interaction by PERMANOVA on Euclidean distances between samples (999 permutations). Fold changes were LB − R2A within each method, and agreement between methods is reported as Pearson *r*. Total protein was compared between methods within each medium by Welch’s *t*-test with Holm correction. For the Flex carbon × nitrogen experiment, fold changes were the carbon and nitrogen main effects, each averaged over the other nutrient, with wells from the same flask treated as correlated (duplicateCorrelation) so that the flask was the replicate. Technical CVs were calculated per protein group from linear-scale intensities across the wells of each flask and summarized for each condition as the median of its three flasks. In both experiments, proteins were called changed at Benjamini–Hochberg adjusted *P* < 0.05 and |log_2_ FC| > 1. For the Flex scale-up plates, CVs were calculated across the wells of each group, and perimeter and interior wells were compared by Welch’s *t*-test.

## Results

### Modified-sequential chemistry and its automation reproduce canonical MPLEx

In canonical MPLEx,^5^ the aqueous phase is drawn off from above, but the organic phase can only be reached through the gelatinous protein disc that forms at the interface (Figure 1 B). Disturbing that disc can strip part or all of the protein layer onto the pipette tip, so the organic draw is the critical step and relies on careful visual monitoring. To avoid this, after the aqueous phase is collected, the protocol dilutes the organic phase with a second methanol aliquot. This lowers the density of the organic phase so that a second centrifugation drives the protein to the bottom of the tube as a firm pellet (Figure 1 C). The organic layer can then be aspirated from above without contacting the protein, stopping at a fixed height above the pellet. We set these aspiration heights empirically, using methylene blue and oil red O to mark the aqueous and organic layers (Figure S2).

To test whether this change preserves biological differences between samples, we extracted *P. putida* grown in two media (LB and R2A) in quadruplicate using three protocols: canonical MPLEx performed by hand (Canonical), the modified OpenMPLEx chemistry performed by hand (Human Sequential), and the same OpenMPLEx chemistry performed on the OT-2 (OT-2 Robot). LB and R2A differ substantially in nutrient content, with R2A formulated as a low-nutrient medium,^24^ and medium composition is known to reshape bacterial growth and proteome allocation,^25^ including in *P. putida*.^26^ The two media therefore provide a biologically meaningful contrast against which to compare manual and automated processing. Our main comparison is the difference each method measures between the two media, rather than absolute yields between methods.

All three methods identified comparable numbers of precursors and protein groups, around 4,200 protein groups per replicate, with replicate variation overlapping across methods and media (Figure 2 A). They also quantified the same proteins: of the 4,336 protein groups quantified in all four replicates of at least one medium, 4,155 were shared by all three methods (Figure S4 A).

**Figure 2:**
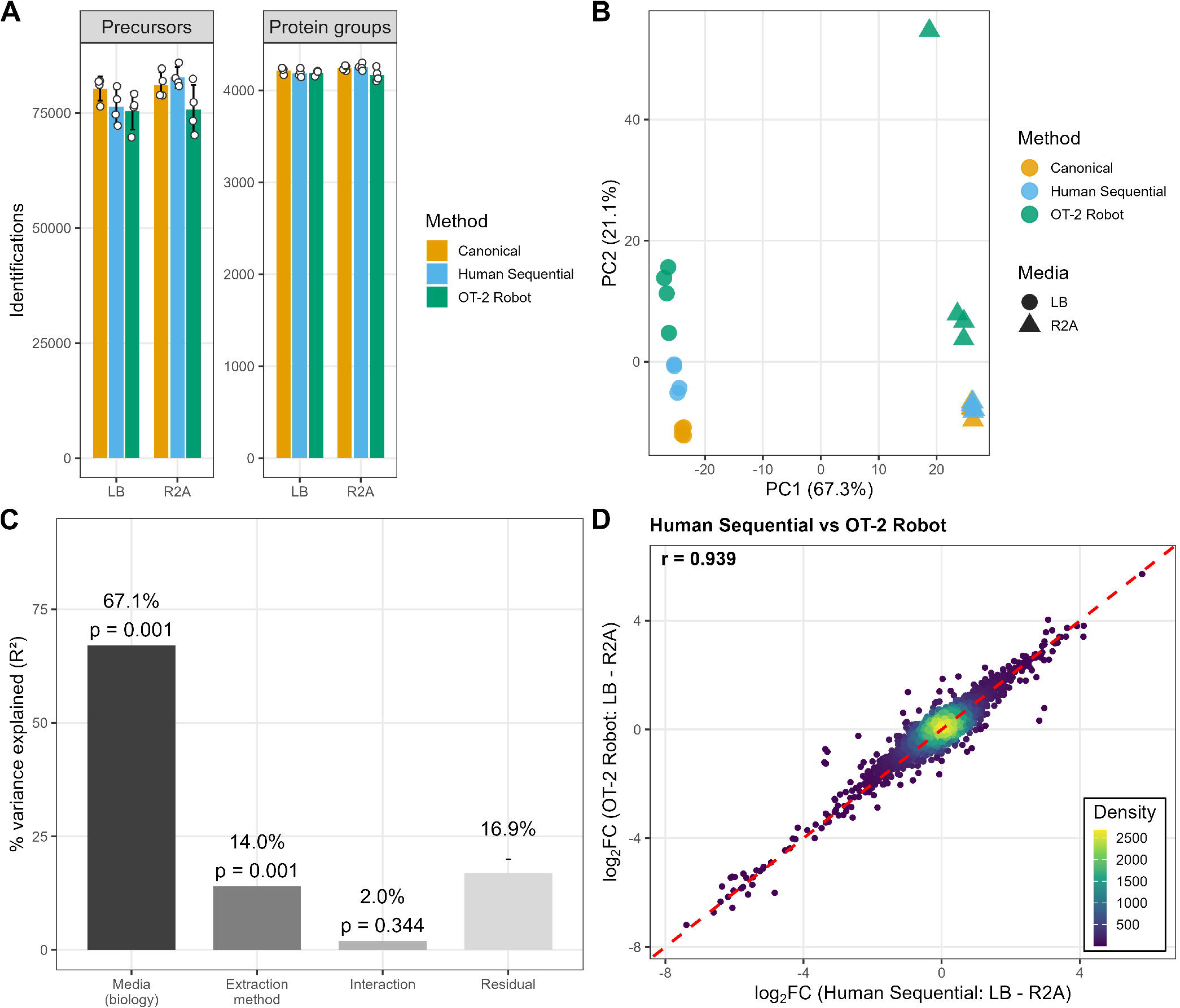
Proof-of-concept validation of OpenMPLEx chemistry on the Opentrons OT-2. (A) Precursor and protein-group identifications per replicate (mean ± SD). (B) PCA of the 3,923 protein groups quantified in every sample; PC1 separates samples by growth media, and PC2 separates OT-2 Robot from the manual methods. One OT-2 R2A replicate appears as a high-PC2 outlier and is retained in all analyses (CV% per group in Figure S5). (C) PERMANOVA variance partition: biology dominates over extraction method, and the media × method interaction is non-significant. (D) Per-protein log₂ fold change (LB − R2A) between OT-2 Robot and Human Sequential extractions; tight scatter around the identity line (red dashed) shows the two methods agree on which proteins are up- and down-regulated.

The OT-2 recovered less total protein on average than either manual method, 69 to 83% of the manual yields, although with four replicates per group no pairwise difference within a medium was significant after correction for multiple testing (Figure S6). Recovery with Human Sequential did not differ significantly from Canonical. We found no evidence that the OT-2 shortfall was selective. Within each medium, the per-protein difference between OT-2 Robot and Human Sequential centered on zero (median log_2_ difference 0.03 in LB and 0.02 in R2A), and 0.5% (LB) and 2.1% (R2A) of protein groups differed by more than 2-fold, against 11.5% between media (Figure S4 B). Even with the lower recovery, there was ample protein for downstream analysis, and all samples were normalized to the same 250 ng load before injection. We attribute the shortfall to sample retained on pipette tips during on-deck mixing. This can be avoided by mixing with a shaker, so that pipettes make no unnecessary contact with samples. We mixed by pipette on the OT-2 because it cannot move plates, and the tubes had to stay on the chiller to maximize phase separation and protein aggregation. More capable robots can move a plate between chiller and shaker, or combine the two in one unit, and we took the first approach on the Flex. LB recovery exceeded R2A in every method.

In principal component analysis of the protein groups quantified in every sample, PC1 (67.3%) separated samples by media, and PC2 (21.1%) separated OT-2 Robot from the manual methods and contained the remaining replicate structure (Figure 2 B). We used PERMANOVA to assign variance to biology or to preparation. Media accounted for 67.1% of the variance (p = 0.001), about five times the 14.0% accounted for by extraction method (p = 0.001), and the media-by-method interaction accounted for 2.0% and was not significant (p = 0.344) (Figure 2 C). Extraction method shifts every condition by a similar amount rather than reshaping the biology, so its effect cancels when LB is compared with R2A. Protein-group CVs were similar for every method-and-media group except OT-2 R2A, where one replicate inflated the group CV (Figure S5). The fault is confined to the proteomics fraction, which places it downstream of extraction, and the TICs suggest an error during Evotip loading. We retained this replicate in every analysis; removing it changes none of the conclusions.

We then tested our primary comparison: whether the methods measure the same differences between conditions. Per-protein log_2_ fold changes between media tracked the identity line in every pairwise comparison (Figure 2 D and Figure S7), indicating that all three methods report similar differences between media. The three-arm design tests each change separately. Canonical against Human Sequential isolates the chemistry change (*r* = 0.948), and Human Sequential against OT-2 Robot isolates automation (*r* = 0.939). Neither change substantially altered the fold changes, and Canonical against OT-2 Robot, which combines both changes, agrees as well (*r* = 0.952). Volcano plots show the same comparison in the form most readers expect, with each method calling a similar number of proteins changed between media (Figure S8). The proteins highlighted in those plots changed in the same direction and by comparable magnitudes in all three methods. Consistent with *P. putida*’s reliance on the Entner–Doudoroff (ED) pathway for glucose catabolism,^27,28^ the ED enzyme Eda and the glucose-uptake porin OprB were more abundant in the glucose-containing, low-nutrient R2A medium. Growth in amino acid-rich LB, where *P. putida* preferentially assimilates amino acids under Crc-mediated catabolite repression,^26,29^ was associated with elevated levels of the arginine catabolic enzyme Ocd.^30^ The largest differences between media were nutritional-stress responses rather than central metabolism: pyoverdine biosynthesis proteins (PvdA, PvdL, PvdD, PvdI, PvdJ) were tens-to over a hundredfold higher in R2A (log_2_FC −5 to −7), consistent with the lower iron availability of R2A relative to LB.

Metabolite and lipid fractions from the same extractions were also analyzed. Because the aqueous phase is collected before the chemistries diverge, metabolite extraction should be unaffected by the modified chemistry, and we measured metabolites primarily as a check. No pooled QC samples or internal standards were included for this fraction, so analytical and extraction variability could not be separated. Fold changes between media agreed across methods (*r* = 0.907–0.930; Figure S9). Unlike the aqueous phase, the organic phase is collected after the methanol addition, so the lipid fraction is the one most likely to be affected by the modified chemistry. Fold changes between media correlated at *r* = 0.856 for Canonical against Human Sequential, *r* = 0.832 for Human Sequential against OT-2 Robot and *r* = 0.758 for Canonical against OT-2 Robot (Figure S10). These samples were acquired in method-blocked run order, so extraction method and run order are confounded in the lipidomics data. SPLASH internal standards were included to monitor analytical precision. Lipid reproducibility, sample correlations and summary statistics are in the Supporting Information (Figure S11, Figure S12, Figure S13 and Table S1).

These results show that the modified chemistry and its automation perform comparably to canonical MPLEx in coverage, in the differences they detect between conditions and in differential abundance, with a small cost in protein recovery on the OT-2. Having established that the approach works in individual tubes, the next step was scaling up to wellplates.

### Semi-automated wellplate scale-up

To increase throughput beyond the tube-based OT-2 protocol, we scaled the workflow to 96-well plates on the Opentrons Flex. Plates are also where automation has the clearest advantage: an operator cannot see the interface in interior wells of a deep-well plate or hold a multichannel pipette at a fixed depth across 80 wells. Each plate holds 80 samples, leaving columns 11 and 12 free for the reduction/alkylation cocktail and bead stock in the SP3 step (S5), BCA standard curves (S2), and optional quality-control samples. All scripts share a single deck layout (Figure 3 A); only the unpowered magnetic block moves between scripts, avoiding slow deck rearrangement. A complete extraction, from sample loading to placing the protein plate in the vacuum centrifuge, takes about 2 h, including two 10-min holds before centrifugation.

**Figure 3:**
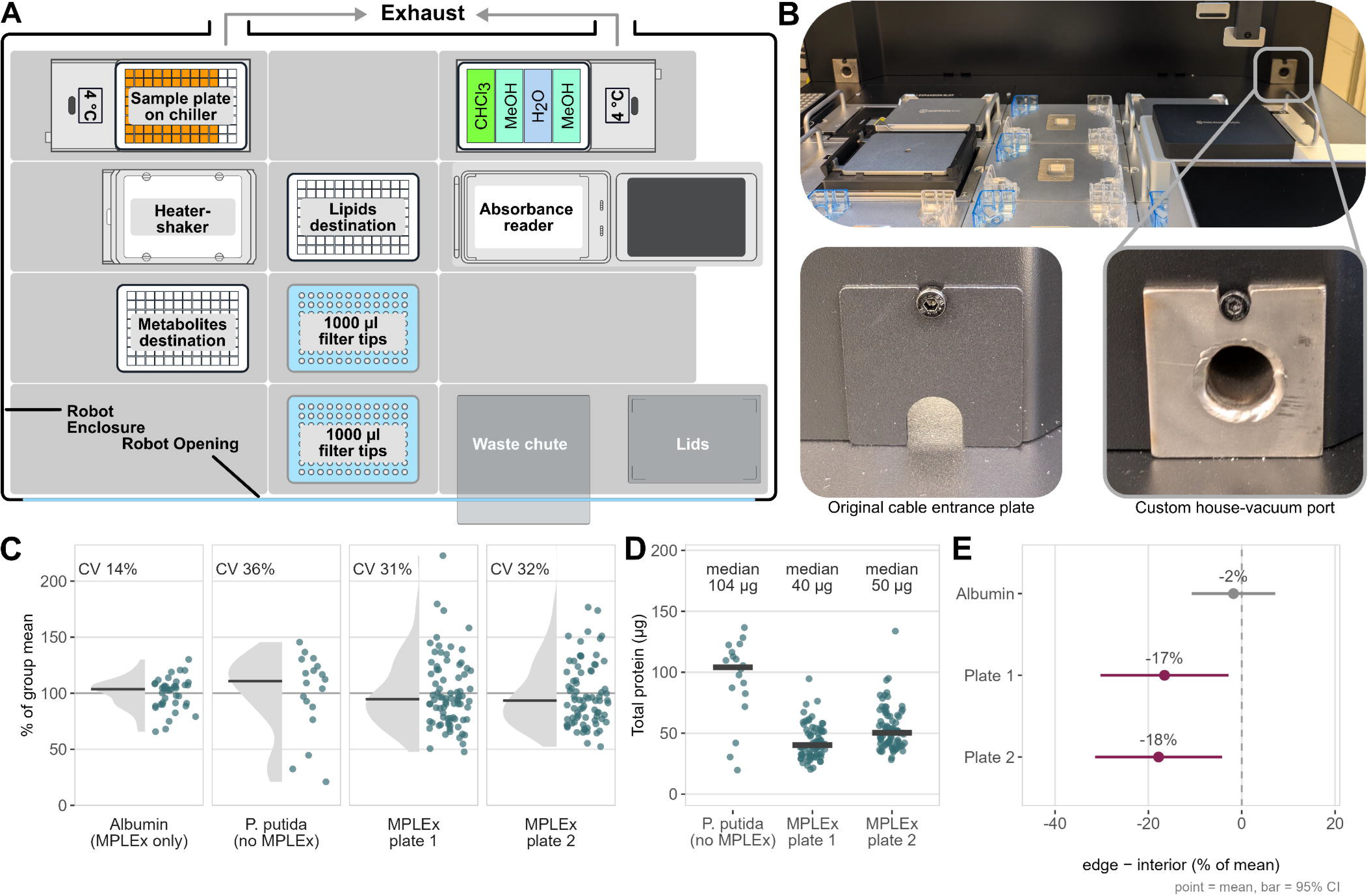
OpenMPLEx scales to the 96-well Opentrons Flex. (A) Flex deck layout for the OpenMPLEx protocol; the modules stay in place across scripts, and only the magnetic block moves. (B) Enclosure ventilation: the original cable-entrance plate (left) was replaced with a custom house-vacuum port (right) to clear chloroform and methanol vapor. (C–E) Reproducibility across two 80-sample *P. putida* plates extracted on different days. (C) Per-well total protein by BCA (% of group mean, CV annotated); *P. putida* MPLEx (31–32%) falls between the same culture lysed without extraction (36%; two empty wells excluded) and albumin carried through MPLEx (14%, the technical floor). (D) Absolute total protein per well (median); MPLEx recovers roughly half the direct-lysate total. (E) Edge effect: perimeter minus interior wells (mean ± 95% CI); no bias in the albumin control, both plates a modest ∼17% deficit.

Institutional safety requirements call for chloroform and methanol to be handled with ventilation. The OT-2 was run in a fume hood, but the Flex does not fit in one, so we vented its manufacturer-supplied enclosure to a house-vacuum line through a custom port replacing the cable-entrance plate (Figure 3 B and Figure S14). Institutional safety and engineering staff measured a constant vacuum pull around the instrument and certified the setup. The protocol further limits exposure: used tips return to their rack so residual chloroform evaporates inside the vented enclosure, 10-min holds before each centrifugation support phase separation and allow solvent vapor to be evacuated before the operator opens the door, and plates are lidded where possible and foil-sealed before centrifugation.

To test reproducibility at this scale, we extracted two 80-sample plates of *P. putida* from a single culture on different days and read per-well total protein by on-deck BCA, taking each well as a percentage of its plate mean. Because every well came from one culture, this variation is technical. The automated extraction varied by 31% and 32% (CV) across the two plates (Figure 3 C). We measured two references to help interpret this range. Albumin carried through the full extraction involves no cells, so its CV of 14% gives a floor for pipetting and the assay. The same cells lysed directly without extraction include cell loading and the assay but no extraction, and varied by 36%. The extraction fell between these references on both days.

The two plates also agreed in absolute terms, with median per-well yields of 40 and 50 µg against 104 µg for the unextracted lysate (Figure 3 D). The lysate value may be an overestimate, because compounds that interfere with the BCA assay, such as reducing sugars, lipids, free amino acids, peptides and chelators, are present in the lysate and are removed by MPLEx.^31^ Perimeter wells gave modestly lower yields than interior wells, a ∼17% edge deficit that was absent in the albumin control (Figure 3 E).

### A designed carbon × nitrogen experiment, end to end

To demonstrate that the entire workflow could retrieve biological information, we constructed a 2 × 2 media comparison with *P. putida* KT2440 grown in minimal medium with either glucose or *p*-coumarate as the carbon source and either NH_4_Cl or NaNO_3_ as the nitrogen source, in three flasks per condition (Figure 4 A). Equal biomass loadings from each flask were randomly distributed across 80 wells of one plate, yielding technical replicates alongside biological replicates (Figure 4 B). One operator completed the full processing, from input plate to acquisition of LC-MS data, in a working week. The workflow took approximately 3 h of total hands-on time and an additional 11 h of automation time, and LC-MS data acquisition took a further 1 h of manual setup and 10 h of unattended acquisition (Table 1).

**Figure 4:**
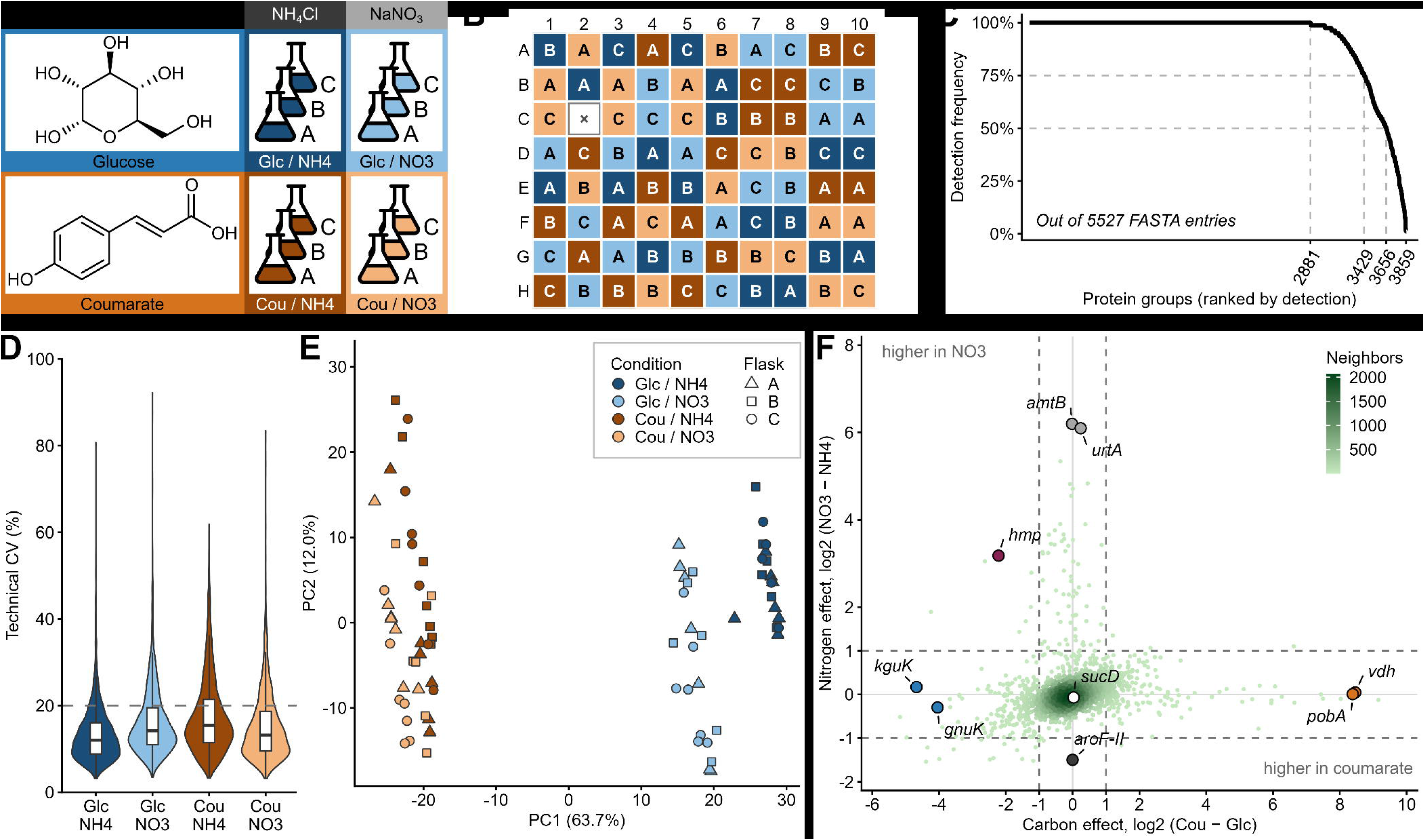
A designed carbon × nitrogen experiment run end to end on the Flex. (A) Growth design, with three flasks (A–C) per condition. (B) Plate layout: fill, condition; letter, flask; ×, failed well C2. (C) Protein groups ranked by detection across the 79 wells. Ticks mark the numbers in all wells, in ≥ 75%, in ≥ 50% and in at least one well. (D) Technical CV per protein group across the wells of a flask, as the median of the condition’s three flasks; dashed line, 20%. (E) PCA of the protein groups quantified in every well; point, well; shape, flask. (F) Carbon effect against nitrogen effect per protein group. Color, number of neighboring proteins; dashed lines, ±1 log_2_. Highlighted are the largest single-nutrient responses, Hmp and an unchanged reference, SucD.

Across the 79 samples, 2,881 protein groups were detected in every sample and 3,859 in at least one (Figure 4 C). One well, C2, failed because of a sample handling error after digestion and upstream of the LC-MS (Figure S15). The median technical CV across the wells of a flask, which captures the whole workflow from extraction to LC-MS, was 12.0–15.4% by condition (Figure 4 D). It fell with intensity, from about 20% for the least abundant protein groups to about 10% for the most abundant (Figure S16). Wells from the same flask correlated at a median *r* of 0.991 (Figure S17). We saw no polymer contamination in the proteomics samples, as indicated by the MS1 maps (Figure S18).

Samples separated by carbon source on PC1 (63.7%) and by nitrogen source within each carbon source (Figure 4 E and Figure S19). To compare the two nutrients directly, we averaged both glucose arms against both coumarate arms for the carbon effect, and both NH_4_ arms against both NO_3_ arms for the nitrogen effect (Figure 4 F). With the flask as the replicate, 496 proteins changed with carbon, 94 with nitrogen and 101 with their interaction (Figure S20 and Figure S21). Most protein groups clustered near the origin, the carbon axis spanned a wider range than the nitrogen axis, and the strongest responses were specific to one nutrient.

The carbon response matches published targeted proteomics of *P. putida* grown with *p*-coumarate or glucose as the carbon source.^32^ In that study, most β-ketoadipate pathway enzymes were higher with *p*-coumarate, most initial glucose catabolism enzymes and all Entner–Doudoroff enzymes were higher with glucose, and few TCA cycle enzymes changed. The same proteins sat at the extremes here, at similar magnitudes. Vanillin dehydrogenase (Vdh) and 4-hydroxybenzoate 3-monooxygenase (PobA), on the route from *p*-coumarate to protocatechuate, were more than 300-fold higher in coumarate. 2-Ketogluconate kinase (KguK) and gluconokinase (GnuK), which phosphorylate the products of periplasmic glucose oxidation, were about 17- to 26-fold higher in glucose. Succinyl-CoA synthetase (SucD) did not change (Figure 4 F).

The nitrogen response was also as expected. Cells grew more slowly with nitrate (Figure S1), and nitrate raised the ammonium transporter AmtB and the urea transporter UrtA by about 60- to 75-fold; scavenging nitrogen through uptake systems is a main response of *P. putida* to nitrogen limitation.^33,34^ Responses higher in NH_4_ were few and small: 76 of the 94 nitrogen-responsive proteins were higher in nitrate, and the largest change in the other direction, AroF-II, was about 1.5 log_2_ units. Hmp, a flavohemoprotein that detoxifies nitric oxide,^35^ changed most with both nutrients: it was higher in nitrate, consistent with nitrosative stress during nitrate assimilation, and higher in glucose. Figure S22 shows the statistical tests for each of these proteins across the four arms. Together, these responses show that the workflow recovered the known differences between conditions.

## Discussion

With a minor change in chemistry, OpenMPLEx avoids drawing the organic phase through the protein layer, and the automated protocol reported differences between media similar to those from the manual protocols (Figure 2). On the Flex, it also worked as a complete workflow: one operator took 80 samples from frozen cell pellets through LC-MS and data analysis in a working week. Sample preparation took about 14 h overall, about 11 h of it on the robot and only about 3 h hands-on, followed by about 10 h of unattended LC-MS (Table 1), whereas preparing the same samples by hand would take days at the bench. The extraction alone takes about 30 min of hands-on time and 2 h on the robot per plate (Table 1), so we estimate that a single operator could run two plates a day, producing all three fractions (protein, lipid, and metabolite) for 160 samples; at that rate the practical bottleneck would be access to the vacuum centrifuge rather than the robot or the operator. During manual aspiration, an overly fast plunger release can disturb the protein pellet, whereas the robot applies the same aspiration speed to every well. Layers are drawn off slowly to avoid disturbing the protein, the second methanol addition is a fast pulse that passes through the protein layer and mixes with the chloroform, and the SP3 script uses slow draws to avoid disturbing the beads.

The median technical CV of 12–15% across the full workflow (Figure 4 D) was low enough to recover the expected carbon and nitrogen responses with three flasks per condition. Automation also makes replication cheap, which matters when a preparation fails. The lost well here cost one of twenty technical replicates in its condition, and the outlier in the OT-2 comparison changed no conclusion. Higher throughput should also compress a study into fewer processing days and batches, which reduces between-batch variance across a cohort. However, the robot cannot detect bubbles, which can sit at the bottom of PCR-plate wells where the pipette aspirates. We therefore recommend checking plates before a transfer and spinning them down briefly between scripts, which also pellets any residual beads. Centrifugation has been reported to reduce the effects of entrapped air and unsettled menisci in low-volume wells.^36^

The platform is also inexpensive and open source, and it requires no integrated centrifuge. The protocols are modular Python scripts that can be adapted to other plates, sample types and downstream assays. We developed them with an LLM coding assistant, and similar tools should lower the programming burden of adapting them. We provide the scripts, labware definitions and print files as is in the Supporting Information, and other laboratories should expect to adjust and test them on their own decks; the partitioning dyes make this easier (Figure S2). The house-vacuum ventilation was simple to build and was certified by our institutional safety and engineering staff (Figure S14). Other laboratories will need to meet their own institutional safety requirements.

The same deck and scripts should carry other chemistries, such as methyl-*tert*-butyl ether extraction,^37^ which places the lipid layer on top, or bead-based multi-omics preparation (BAMM),^38^ which needs no phase separation, and we are pursuing these on the same platform. MPLEx remains a practical choice wherever existing data and methods are built on it or where pathogen inactivation is needed,^11^ although we did not test inactivation with the sequential chemistry.

## Conclusions

OpenMPLEx brings MPLEx to low-cost, open-source liquid handlers, using a minor change in chemistry to circumvent a technically challenging engineering problem. The sequential chemistry and its automation reported differences between media similar to those from canonical manual MPLEx, and the workflow scaled to 96-well plates. In a full run, one operator took 80 samples from cell pellets to LC-MS data in a working week with approximately 3 h of hands-on time, and the data recovered the expected carbon and nitrogen responses. We provide the scripts, labware definitions and print files so that other laboratories can adapt the workflow to their own samples and downstream analyses.

## Supporting information

scripts and labware

Proteomics Data

Lipids and Metabolite data

## Associated Content

The Supporting Information is available free of charge:

- Supplemental methods for metabolomics and lipidomics, Figures S1–S22 and Table S1
- OpenMPLEx_SI_scripts_and_labware.zip: protocol scripts, custom labware definitions, STEP files for 3D-printed labware and the preparation workbook
- OpenMPLEx_SI_protein_group_matrices.xlsx: protein group intensities and sample design for both proteomics experiments
- OpenMPLEx_SI_lipid_metabolite_feature_matrices.xlsx: xcms feature intensities and sample design for the lipidomics and metabolomics fractions of the OT-2 extraction comparison
- OpenMPLEx_SI_BCA_data.xlsx: BCA measurements per well for Figure 3, total protein for the OT-2 extraction comparison, and protein and peptide recovered per well on the carbon × nitrogen plate

## Data Availability

The proteomics, lipidomics and metabolomics mass spectrometry data, with the DIA-NN search results and xcms feature-detection outputs, have been deposited in MassIVE under accession MSV000103995 and are registered with ProteomeXchange under accession PXD087323. Protein group matrices, lipidomics and metabolomics feature matrices, BCA measurements, protocol scripts, labware definitions and STEP files are provided in the Supporting Information; the scripts are available under the MIT License.

## Author Contributions

D.R.H., K.D., and M.A.G. conceived the study. D.R.H. developed the methodology and, with J.B.T., wrote the automation software; J.B.T. also assisted with robotic sample processing, and O.S. provided advice on automation. E.L.B. performed all bacterial culturing and selected the media for the biological experiments. K.L.T. performed sample preparation for the manual arm of the methods comparison. D.R.H. acquired the proteomics data, J.G.E. the lipidomics data, and P.M.L. the metabolomics data. D.R.H. analyzed and visualized the data. P.P. and T.L. provided resources and instrumentation support. D.R.H. and M.A.G. administered the project. D.R.H. wrote the original draft, and all authors reviewed and edited the manuscript.

## Notes

The authors declare no competing financial interest.

## Acknowledgments

The authors thank Matthew Gaffrey and Sarah Fansler for their assistance in assessing ventilation of the Opentrons Flex robot and securing institutional approval for its use, EMSL Crafts for fabricating the custom ventilation adaptors, Adam Ryan of the EMSL IDAL makerspace for assistance with 3D printing of custom labware, and Matthew Monroe for assistance with data upload to MassIVE. Assistance with robotic automation scripts, data analysis and writing was provided by Claude Opus 5.5 (Anthropic); all AI-assisted code and text were reviewed and validated by the authors, who tested the code and take full responsibility for the work. This research was performed on project awards (DOI: 10.46936/intm.proj.2025.61822/60013815, awarded to D.R.H. and K.D.; DOI: 10.46936/intm.proj.2025.61864/60015450, awarded to M.A.G. and O.S.) under the auspices of the Science & Technology portfolio from the Environmental Molecular Sciences Laboratory, a DOE Office of Science User Facility sponsored by the Biological and Environmental Research program under Contract No. DE-AC05-76RL01830.

## References

(1) Arıkan, M.; Muth, T. Integrated Multi-Omics Analyses of Microbial Communities: A Review of the Current State and Future Directions. Molecular Omics 2023, 19 (8), 607–623. 10.1039/D3MO00089C.

(2) Hasin, Y.; Seldin, M.; Lusis, A. Multi-Omics Approaches to Disease. Genome Biology 2017, 18 (1), 83. 10.1186/s13059-017-1215-1.

(3) Guzman, U. H.; Martinez-Val, A.; Ye, Z.; Damoc, E.; Arrey, T. N.; Pashkova, A.; Renuse, S.; Denisov, E.; Petzoldt, J.; Peterson, A. C.; Harking, F.; Østergaard, O.; Rydbirk, R.; Aznar, S.; Stewart, H.; Xuan, Y.; Hermanson, D.; Horning, S.; Hock, C.; Makarov, A.; Zabrouskov, V.; Olsen, J. V. Ultra-Fast Label-Free Quantification and Comprehensive Proteome Coverage with Narrow-Window Data-Independent Acquisition. Nature Biotechnology 2024, 1–12. 10.1038/s41587-023-02099-7.

(4) Billoir, E.; Navratil, V.; Blaise, B. J. Sample Size Calculation in Metabolic Phenotyping Studies. Briefings in Bioinformatics 2015, 16 (5), 813–819. 10.1093/bib/bbu052.

(5) Nakayasu, E. S.; Nicora, C. D.; Sims, A. C.; Burnum-Johnson, K. E.; Kim, Y.-M.; Kyle, J. E.; Matzke, M. M.; Shukla, A. K.; Chu, R. K.; Schepmoes, A. A.; Jacobs, J. M.; Baric, R. S.; Webb-Robertson, B.-J.; Smith, R. D.; Metz, T. O. MPLEx: A Robust and Universal Protocol for Single-Sample Integrative Proteomic, Metabolomic, and Lipidomic Analyses. mSystems 2016, 1 (3), 10.1128/msystems.00043–16. 10.1128/msystems.00043-16.

(6) Duchesneau, K.; Aldeguer-Riquelme, B.; Petro, C.; Makke, G.; Green, M.; Tfaily, M.; Wilson, R.; Roth, S. W.; Johnston, E. R.; Kluber, L. A.; Schadt, C. W.; Trejo, J. B.; Callister, S. J.; Purvine, S. O.; Chanton, J. P.; Hanson, P. J.; Tringe, S.; Eloe-Fadrosh, E.; Glavina Del Rio, T.; Konstantinidis, K. T.; Kostka, J. E. Northern Peatland Microbial Communities Exhibit Resistance to Warming and Acquire Electron Acceptors from Soil Organic Matter. Nature Communications 2025, 16 (1), 6869. 10.1038/s41467-025-61664-7.

(7) Del Cerro, C.; Erickson, E.; Dong, T.; Wong, A. R.; Eder, E. K.; Purvine, S. O.; Mitchell, H. D.; Weitz, K. K.; Markillie, L. M.; Burnet, M. C.; Hoyt, D. W.; Chu, R. K.; Cheng, J.-F.; Ramirez, K. J.; Katahira, R.; Xiong, W.; Himmel, M. E.; Subramanian, V.; Linger, J. G.; Salvachúa, D. Intracellular Pathways for Lignin Catabolism in White-Rot Fungi. Proceedings of the National Academy of Sciences 2021, 118 (9), e2017381118. 10.1073/pnas.2017381118.

(8) Kim, J.; Coradetti, S. T.; Kim, Y.-M.; Gao, Y.; Yaegashi, J.; Zucker, J. D.; Munoz, N.; Zink, E. M.; Burnum-Johnson, K. E.; Baker, S. E.; Simmons, B. A.; Skerker, J. M.; Gladden, J. M.; Magnuson, J. K. Multi-Omics Driven Metabolic Network Reconstruction and Analysis of Lignocellulosic Carbon Utilization in Rhodosporidium Toruloides. Frontiers in Bioengineering and Biotechnology 2021, 8. 10.3389/fbioe.2020.612832.

(9) Berrios, L.; Bogar, G. D.; Bogar, L. M.; Venturini, A. M.; Willing, C. E.; Del Rio, A.; Ansell, T. B.; Zemaitis, K.; Velickovic, M.; Velickovic, D.; Pellitier, P. T.; Yeam, J.; Hutchinson, C.; Bloodsworth, K.; Lipton, M. S.; Peay, K. G. Ectomycorrhizal Fungi Alter Soil Food Webs and the Functional Potential of Bacterial Communities. mSystems 2024, 9 (6), e0036924. 10.1128/msystems.00369-24.

(10) Gregory, D. J.; Han, F.; Li, P.; Gritsenko, M. A.; Kyle, J.; Riley, F. E.; Chavez, D.; Yotova, V.; Sindeaux, R. H. M.; Hawash, M. B. F.; Xu, F.; Hung, L.-Y.; Hayden, D. L.; Tompkins, R. G.; Lanford, R. E.; Kobzik, L.; Hellman, J.; Jacobs, J. M.; Barreiro, L. B.; Xiao, W.; Warren, H. S. Molecular Profiles of Blood from Numerous Species That Differ in Sensitivity to Acute Inflammation. Molecular Medicine 2024, 30 (1), 280. 10.1186/s10020-024-01052-x.

(11) Burnum-Johnson, K. E.; Kyle, J. E.; Eisfeld, A. J.; Casey, C. P.; Stratton, K. G.; Gonzalez, J. F.; Habyarimana, F.; Negretti, N. M.; Sims, A. C.; Chauhan, S.; Thackray, L. B.; Halfmann, P. J.; Walters, K. B.; Kim, Y.-M.; Zink, E. M.; Nicora, C. D.; Weitz, K. K.; Webb-Robertson, B.-J. M.; Nakayasu, E. S.; Ahmer, B.; Konkel, M. E.; Motin, V.; Baric, R. S.; Diamond, M. S.; Kawaoka, Y.; Waters, K. M.; Smith, R. D.; Metz, T. O. MPLEx: A Method for Simultaneous Pathogen Inactivation and Extraction of Samples for Multi-Omics Profiling. The Analyst 2017, 142 (3), 442–448. 10.1039/C6AN02486F.

(12) Nicora, C. D.; Burnum-Johnson, K. E.; Nakayasu, E. S.; Casey, C. P.; White, R. A.; Roy Chowdhury, T.; Kyle, J. E.; Kim, Y.-M.; Smith, R. D.; Metz, T. O.; Jansson, J. K.; Baker, E. S. The MPLEx Protocol for Multi-omic Analyses of Soil Samples. Journal of Visualized Experiments: JoVE 2018, No. 135, 57343. 10.3791/57343.

(13) Chen, Y.; Guenther, J. M.; Gin, J. W.; Chan, L. J. G.; Costello, Z.; Ogorzalek, T. L.; Tran, H. M.; Blake-Hedges, J. M.; Keasling, J. D.; Adams, P. D.; García Martín, H.; Hillson, N. J.; Petzold, C. J. Automated “Cells-To-Peptides” Sample Preparation Workflow for High-Throughput, Quantitative Proteomic Assays of Microbes. Journal of Proteome Research 2019, 18 (10), 3752–3761. 10.1021/acs.jproteome.9b00455.

(14) Moukarzel, G.; Wang, Y.; Xin, W.; Hofmann, C.; Joshi, A.; Loughney, J. W.; Bowman, A. Automation of Biochemical Assays Using an Open-Sourced, Inexpensive Robotic Liquid Handler. SLAS Technology 2024, 29 (6), 100205. 10.1016/j.slast.2024.100205.

(15) Kverneland, A. H.; Harking, F.; Vej-Nielsen, J. M.; Huusfeldt, M.; Bekker-Jensen, D. B.; Svane, I. M.; Bache, N.; Olsen, J. V. Fully Automated Workflow for Integrated Sample Digestion and Evotip Loading Enabling High-Throughput Clinical Proteomics. Molecular & Cellular Proteomics 2024, 23 (7). 10.1016/j.mcpro.2024.100790.

(16) Hughes, C. S.; Moggridge, S.; Müller, T.; Sorensen, P. H.; Morin, G. B.; Krijgsveld, J. Single-Pot, Solid-Phase-Enhanced Sample Preparation for Proteomics Experiments. Nature Protocols 2019, 14 (1), 68–85. 10.1038/s41596-018-0082-x.

(17) Kulak, N. A.; Pichler, G.; Paron, I.; Nagaraj, N.; Mann, M. Minimal, Encapsulated Proteomic-Sample Processing Applied to Copy-Number Estimation in Eukaryotic Cells. Nature Methods 2014, 11 (3), 319–324. 10.1038/nmeth.2834.

(18) Bache, N.; Geyer, P. E.; Bekker-Jensen, D. B.; Hoerning, O.; Falkenby, L.; Treit, P. V.; Doll, S.; Paron, I.; Müller, J. B.; Meier, F.; Olsen, J. V.; Vorm, O.; Mann, M. A Novel LC System Embeds Analytes in Pre-formed Gradients for Rapid, Ultra-robust Proteomics*. Molecular & Cellular Proteomics 2018, 17 (11), 2284–2296. 10.1074/mcp.TIR118.000853.

(19) Demichev, V.; Messner, C. B.; Vernardis, S. I.; Lilley, K. S.; Ralser, M. DIA-NN: Neural Networks and Interference Correction Enable Deep Proteome Coverage in High Throughput. Nature Methods 2020, 17 (1), 41–44. 10.1038/s41592-019-0638-x.

(20) Mellacheruvu, D.; Wright, Z.; Couzens, A. L.; Lambert, J.-P.; St-Denis, N.; Li, T.; Miteva, Y. V.; Hauri, S.; Sardiu, M. E.; Low, T. Y.; Halim, V. A.; Bagshaw, R. D.; Hubner, N. C.; al-Hakim, A.; Bouchard, A.; Faubert, D.; Fermin, D.; Dunham, W. H.; Goudreault, M.; Lin, Z.-Y.; Badillo, B. G.; Pawson, T.; Durocher, D.; Coulombe, B.; Aebersold, R.; Superti-Furga, G.; Colinge, J.; Heck, A. J. R.; Choi, H.; Gstaiger, M.; Mohammed, S.; Cristea, I. M.; Bennett, K. L.; Washburn, M. P.; Raught, B.; Ewing, R. M.; Gingras, A.-C.; Nesvizhskii, A. I. The CRAPome: A Contaminant Repository for Affinity Purification Mass Spectrometry Data. Nature methods 2013, 10 (8), 730–736. 10.1038/nmeth.2557.

(21) Demichev, V. DIA-NN: Documentation and Source Repository, 2026. https://github.com/vdemichev/DiaNN (accessed 2026-11-05).

(22) Ritchie, M. E.; Phipson, B.; Wu, D.; Hu, Y.; Law, C. W.; Shi, W.; Smyth, G. K. limma Powers Differential Expression Analyses for RNA-Sequencing and Microarray Studies. Nucleic Acids Research 2015, 43 (7), e47. 10.1093/nar/gkv007.

(23) Oksanen, J.; Simpson, G. L.; Blanchet, F. G.; Kindt, R.; Legendre, P.; others. Vegan: Community Ecology Package; 2026. 10.32614/CRAN.package.vegan.

(24) Reasoner, D. J.; Geldreich, E. E. A New Medium for the Enumeration and Subculture of Bacteria from Potable Water. Applied and Environmental Microbiology 1985, 49 (1), 1–7. 10.1128/aem.49.1.1-7.1985.

(25) Schmidt, A.; Kochanowski, K.; Vedelaar, S.; Ahrné, E.; Volkmer, B.; Callipo, L.; Knoops, K.; Bauer, M.; Aebersold, R.; Heinemann, M. The Quantitative and Condition-Dependent Escherichia Coli Proteome. Nature Biotechnology 2016, 34 (1), 104–110. 10.1038/nbt.3418.

(26) Moreno, R.; Martínez-Gomariz, M.; Yuste, L.; Gil, C.; Rojo, F. The Pseudomonas Putida Crc Global Regulator Controls the Hierarchical Assimilation of Amino Acids in a Complete Medium: Evidence from Proteomic and Genomic Analyses. Proteomics 2009, 9 (11), 2910–2928. 10.1002/pmic.200800918.

(27) Castillo, T. del; Ramos, J. L.; Rodríguez-Herva, J. J.; Fuhrer, T.; Sauer, U.; Duque, E. Convergent Peripheral Pathways Catalyze Initial Glucose Catabolism in Pseudomonas Putida: Genomic and Flux Analysis. Journal of Bacteriology 2007, 189 (14), 5142–5152. 10.1128/JB.00203-07.

(28) Nikel, P. I.; Chavarría, M.; Fuhrer, T.; Sauer, U.; Lorenzo, V. de. Pseudomonas Putida KT2440 Strain Metabolizes Glucose Through a Cycle Formed by Enzymes of the Entner-Doudoroff, Embden-Meyerhof-Parnas, and Pentose Phosphate Pathways. The Journal of Biological Chemistry 2015, 290 (43), 25920–25932. 10.1074/jbc.M115.687749.

(29) Molina, L.; Rosa, R. L.; Nogales, J.; Rojo, F. Pseudomonas Putida KT2440 Metabolism Undergoes Sequential Modifications During Exponential Growth in a Complete Medium as Compounds Are Gradually Consumed. Environmental Microbiology 2019, 21 (7), 2375–2390. 10.1111/1462-2920.14622.

(30) Lu, C.-D. Pathways and Regulation of Bacterial Arginine Metabolism and Perspectives for Obtaining Arginine Overproducing Strains. Applied Microbiology and Biotechnology 2006, 70 (3), 261–272. 10.1007/s00253-005-0308-z.

(31) Reichelt, W. N.; Waldschitz, D.; Herwig, C.; Neutsch, L. Bioprocess Monitoring: Minimizing Sample Matrix Effects for Total Protein Quantification with Bicinchoninic Acid Assay. Journal of Industrial Microbiology & Biotechnology 2016, 43 (9), 1271–1280. 10.1007/s10295-016-1796-9.

(32) Gao, Y.; Fillmore, T. L.; Munoz, N.; Bentley, G. J.; Johnson, C. W.; Kim, J.; Meadows, J. A.; Zucker, J. D.; Burnet, M. C.; Lipton, A. K.; Bilbao, A.; Orton, D. J.; Kim, Y.-M.; Moore, R. J.; Robinson, E. W.; Baker, S. E.; Webb-Robertson, B.-J. M.; Guss, A. M.; Gladden, J. M.; Beckham, G. T.; Magnuson, J. K.; Burnum-Johnson, K. E. High-Throughput Large-Scale Targeted Proteomics Assays for Quantifying Pathway Proteins in Pseudomonas Putida KT2440. Frontiers in Bioengineering and Biotechnology 2020, 8. 10.3389/fbioe.2020.603488.

(33) Hervás, A. B.; Canosa, I.; Santero, E. Transcriptome Analysis of Pseudomonas Putida in Response to Nitrogen Availability. Journal of Bacteriology 2008, 190 (1), 416–420. 10.1128/JB.01230-07.

(34) Hervás, A. B.; Canosa, I.; Little, R.; Dixon, R.; Santero, E. NtrC-Dependent Regulatory Network for Nitrogen Assimilation in Pseudomonas Putida. Journal of Bacteriology 2009, 191 (19), 6123–6135. 10.1128/JB.00744-09.

(35) Frey, A. D.; Farrés, J.; Bollinger, C. J. T.; Kallio, P. T. Bacterial Hemoglobins and Flavohemoglobins for Alleviation of Nitrosative Stress in Escherichia Coli. Applied and Environmental Microbiology 2002, 68 (10), 4835–4840. 10.1128/AEM.68.10.4835-4840.2002.

(36) Petersen, J.; Nguyen, J. Comparison of Absorbance and Fluorescence Methods for Determining Liquid Dispensing Precision. SLAS Technology 2005, 10 (2), 82–87. 10.1016/j.jala.2005.01.009.

(37) Matyash, V.; Liebisch, G.; Kurzchalia, T. V.; Shevchenko, A.; Schwudke, D. Lipid Extraction by Methyl-Tert-Butyl Ether for High-Throughput Lipidomics. Journal of Lipid Research 2008, 49 (5), 1137–1146. 10.1194/jlr.D700041-JLR200.

(38) Muehlbauer, L. K.; Jen, A.; Zhu, Y.; He, Y.; Shishkova, E.; Overmyer, K. A.; Coon, J. J. Rapid Multi-Omics Sample Preparation for Mass Spectrometry. Analytical Chemistry 2023, 95 (2), 659–667. 10.1021/acs.analchem.2c02042.

