## Supplementary Information for "An open-source semi-automated workflow for multi-omics sample extraction"

#### Supplemental Methods

##### Metabolomics LC-MS/MS analysis

Aqueous (metabolomics) fractions were dried in a SpeedVac and reconstituted in 150 µL of 80:20 methanol:water. Samples were vortexed and sonicated to redissolve, then centrifuged at 10,000 × g for 5 min at 4 °C to pellet any insoluble material. Supernatants were transferred to LC-MS vials, avoiding any pellet.

Polar metabolite extracts were analyzed on a Thermo Vanquish Flex UHPLC system coupled to an Orbitrap Eclipse Tribrid mass spectrometer (Thermo Fisher Scientific) equipped with a heated electrospray ionization (HESI) source, operated in positive ion mode. Reversed-phase separation was performed on a Hypersil GOLD column (2.1 × 150 mm, 3 µm; Thermo Fisher Scientific) at 40 °C using 0.1% formic acid in water (mobile phase A) and 0.1% formic acid in acetonitrile (mobile phase B). The gradient (400 µL/min unless noted) began isocratic at 10% B for 2 min, increased linearly to 90% B from 2–11 min, was held at 90% B from 11–12 min, then maintained at 90% B for an additional 0.5 min at 500 µL/min (12–12.5 min). The composition was returned to 10% B from 12.5–13.5 min and held until 14 min at 500 µL/min, after which the flow rate was reduced to 400 µL/min and the column was re-equilibrated at 10% B for the remainder of the 16 min run.

HESI source parameters were: spray voltage 3.5 kV, ion transfer tube 320 °C, vaporizer 250 °C, sheath/auxiliary/sweep gas 50/10/1 (arbitrary units), and RF lens 70%. MS1 spectra were acquired in the Orbitrap at 120,000 resolution (FWHM at m/z 200) over m/z 67–1,000, with a normalized AGC target of 100% ($4 \times 10^5$) and a maximum injection time of 100 ms. Data-dependent MS2 spectra were acquired on the top 12 most intense precursors per cycle (minimum intensity threshold $2 \times 10^4$) with a 1.6 m/z isolation window and dynamic exclusion of 3 s (±5 ppm, isotopes excluded). HCD spectra were collected in the Orbitrap at 15,000 resolution using stepped normalized collision energies of 20%, 40%, and 60% (AGC $5 \times 10^4$, maximum injection time 50 ms), and CID spectra were collected in the linear ion trap at a fixed normalized collision energy of 30% with a 1.5 m/z isolation window (AGC $1 \times 10^4$, maximum injection time 35 ms).

**Lipidomics LC-MS/MS analysis**

Organic (lipidomics) fractions were dried in a SpeedVac and reconstituted in 300 μL of 2:1 chloroform:methanol. Samples were vortexed and sonicated to redissolve, then centrifuged at 10,000 × g for 5 min at 4 °C. Supernatants were transferred to Waters LC-MS vials, avoiding any pellet, and stored at -80 °C.

Prior to LC-MS analysis, samples were dried in vacuo for 45 min, and reconstituted in 20 μL ethanol-stabilized chloroform followed by 180 μL methanol. A pooled QC sample was prepared by combining 20 μL aliquots from each sample. Reversed-phase separation was performed on a Waters Acquity H-Class CL system using a Waters Acquity CSH C18 column (3.0 × 150 mm, 1.7 μm) with a matched VanGuard pre-column held at 50 °C, with a 5 μL injection volume and a flow rate of 0.3 mL/min. Mobile phase A was 10 mM ammonium acetate in 60:40 water:acetonitrile, and mobile phase B was 10 mM ammonium acetate in 90:10 isopropanol:acetonitrile. The 27-min gradient (in % B) was: 40% at 0 min; ramped to 62% by 1 min, 66% by 4 min, 78% by 9 min, 87% by 11 min, and 99% by 15 min; held at 99% from 15–21 min; returned to 40% by 21.1 min, briefly ramped to 99% (21.1–21.6 min) and back to 40% (21.6–21.7 min); then held at 40% until 27 min. The pooled QC was injected before sample acquisition and between sample blocks to monitor mass accuracy, retention time, and signal stability. Avanti Brain Total Lipid Extract (0.25 mg/mL in methanol) was injected before and after the sample queue as an additional system-suitability standard, and 10:90 methanol:ethanol-stabilized chloroform solvent blanks were injected before, between, and after sample blocks to assess contamination and carryover.

The LC was coupled to an Orbitrap Fusion Lumos Tribrid mass spectrometer (Thermo Fisher Scientific) equipped with a heated electrospray ionization (HESI) source operated in positive ion mode. Source parameters were: spray voltage 3.5 kV, ion transfer tube 300 °C, vaporizer 350 °C, sheath/auxiliary/sweep gas 40/5/1 (arbitrary units), and RF lens 30%. MS1 spectra were acquired in the Orbitrap at 120,000 resolution (FWHM at m/z 200) over m/z 200–1,800, with a normalized AGC target of 100% ($4 \times 10^5$) and a maximum injection time of 50 ms. Data-dependent MS2 spectra were acquired in cycle-time mode (0.7 s) using monoisotopic precursor selection (MIPS, small-molecule mode); precursors were selected above an absolute intensity threshold of $2.5 \times 10^4$ (or 20% relative intensity) and excluded after two selections for 4 s (±10 ppm, isotopes excluded). Each precursor was fragmented in two scan events: HCD spectra were collected in the Orbitrap at 7,500 resolution using stepped normalized collision energies of 25%, 30%, and 35% (1 m/z isolation window, first mass m/z 75, 50 ms maximum injection time, normalized AGC 200% [$1 \times 10^5$]); CID spectra were collected in the linear ion trap at a fixed normalized collision energy of 38% (1 m/z isolation window, activation time 10 ms, activation Q 0.18, 35 ms maximum injection time, normalized AGC 200% [$2 \times 10^4$]).

**Metabolomics and lipidomics data analysis**

Raw `.raw` files were converted to mzML (centroided MS2; profile MS1 for lipidomics, centroided MS1 for metabolomics) using ProteoWizard MSConvert. Feature detection was performed in R 4.5.1 with the Bioconductor `xcms` package (v4.8.0). Peaks were detected with the CentWave algorithm (5 ppm; peakwidth 5–30 s for lipidomics, 5–20 s for metabolomics; $S/N \geq 10$ for lipidomics, $\geq 15$ for metabolomics; prefilter 3 scans at $1 \times 10^3$). Correspondence used peak-density grouping (binSize 0.005 Da; minFraction 0.5 for lipidomics, 0.75 for metabolomics) followed by loess retention-time alignment (PeakGroupsParam, span 0.4, minFraction 0.85) and a final re-grouping pass. Missing values were imputed with `fillChromPeaks(ChromPeakAreaParam())`.

For lipidomics, 6 column blank (`Blank_Co`) injections were used for blank subtraction: a feature was retained only if its maximum group median (across the six method × media groups) exceeded 3 × the median across blank injections. Method blanks contained the SPLASH internal-standard mixture and were therefore excluded from blank subtraction. For metabolomics, the four blank injections were not used in the analysis, and no blank subtraction was performed.

To distinguish features carrying group-structured variance from features dominated by replicate noise, a within-group F-statistic was computed per feature on $\log_2$-transformed intensities: $F_{\text{within}} = \hat{\sigma}^2_{\text{overall}} / \hat{\sigma}^2_{\text{within-group}}$, where the within-group variance was pooled across the six method × media groups. Features with $F_{\text{within}} > 3$ were retained for multivariate and fold-change analysis ("signal features"). This filter selects features whose variance is structured at the group level rather than scattered randomly within replicates; it does *not* attribute that variance specifically to biology. Method-specific extraction artifacts, run-order effects, and media carryover would also pass the threshold and are not partitioned out.

MS2 library searching against the MoNA-LipidBlast and MoNA LC-MS/MS Positive Mode databases was attempted with `MetaboAnnotation::matchSpectra()` (5 ppm precursor tolerance, 20 ppm fragment tolerance, normalized dot-product ≥ 0.7). For lipidomics, two features yielded high-confidence (Schymanski Level 2) MS/MS matches across the entire feature set; the structural ceiling on in-silico-vs-experimental matching for DDA Lumos data led us to omit these from the figures and rely on m/z-based matching against the Avanti UltimateSPLASH ONE deuterated lipid standards (69 / 69 detected within 5 ppm) for class-level anchors. Metabolomics annotation against MoNA yielded too few hits to report and is not shown.

Feature areas were $\log_2$-transformed and median-centered per sample. Differential features (LB vs R2A within each method) were tested with per-feature Welch *t*-tests and Benjamini–Hochberg FDR correction; significance was defined as adjusted $p < 0.05$ with $|\log_2 \text{FC}| > 1$. Feature matrices are provided in `OpenMPLEx_SI_lipid_metabolite_feature_matrices.xlsx`. The full xcms outputs, with software versions and library checksums for the lipidomics annotation, are deposited in MassIVE.

### Supplemental Figures

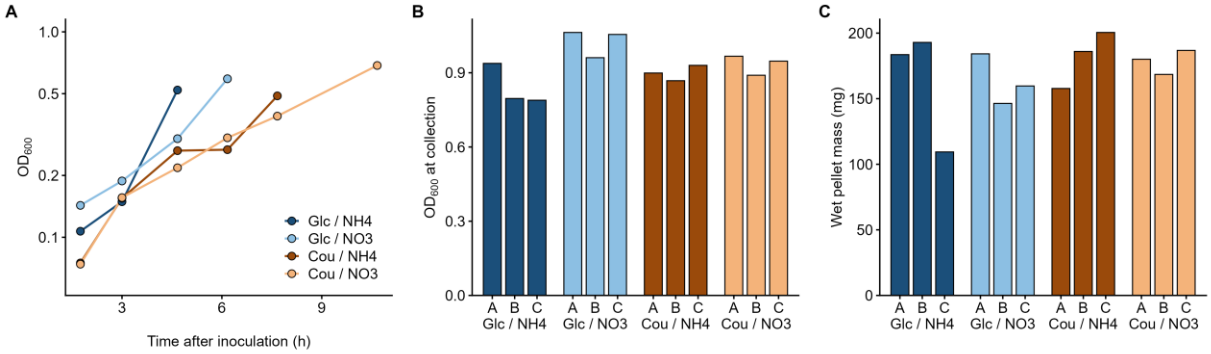


**Figure S1: Growth of the carbon × nitrogen cultures.** (A) $OD_{600}$ during growth, with one reading per condition at each time point, taken from different flasks of that condition. All conditions were inoculated at the same dilution from the acclimation flask. (B) $OD_{600}$ at collection for each flask. (C) Wet pellet mass recovered from each flask.

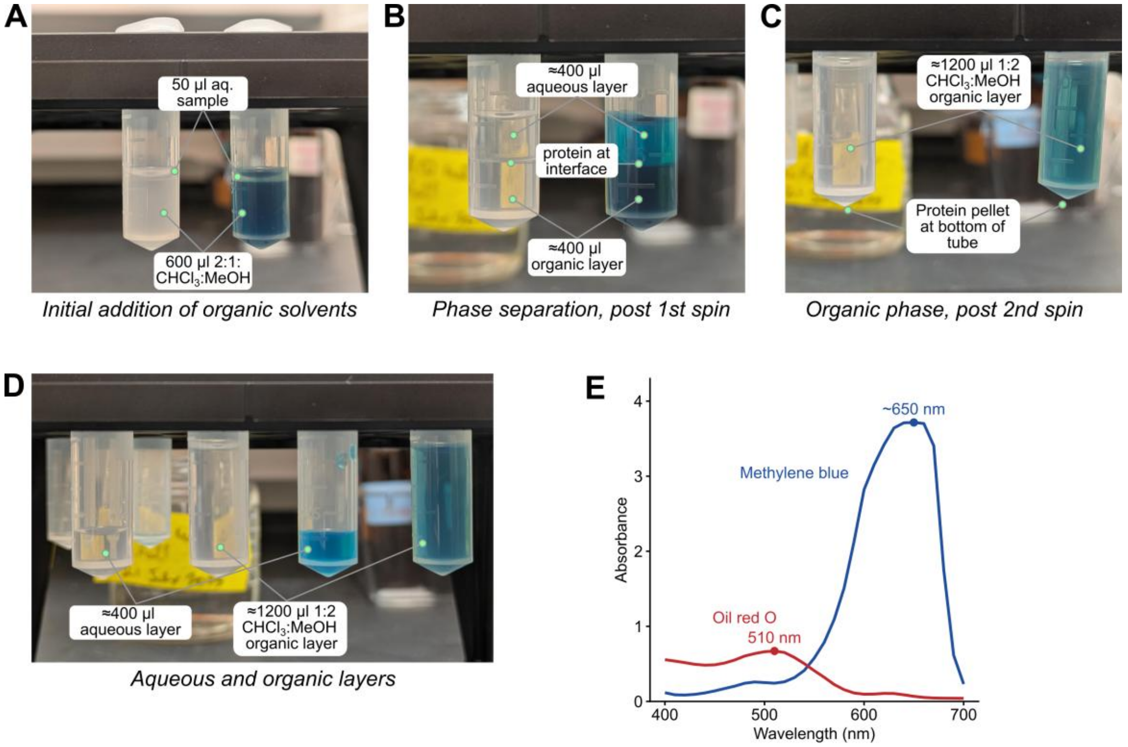


**Figure S2: Partitioning dyes for setting layer heights.** Visualization of phase separations throughout OpenMPLEx extraction. (A) Initial setup with a 50 µL aqueous sample. (B) Phase separation after the first centrifugation. (C) Homogeneous organic phase after methanol (MeOH) addition and the second centrifugation. (D) Collected aqueous and organic layers. (E) Visible absorbance spectra of the two dyes in ethanol (Synergy H1 plate reader, 10 nm steps, 5 µL of each dye stock). Oil red O peaks at 510 nm. Methylene blue peaks near 650 nm, close to the reader's upper limit, so its apex is approximate. The dyes mark the aqueous and organic layers by eye, which makes layer heights easier to set and troubleshoot. Because their absorbance peaks are well separated, a plate reader could also follow each dye in the collected fractions, but we did not find this necessary.

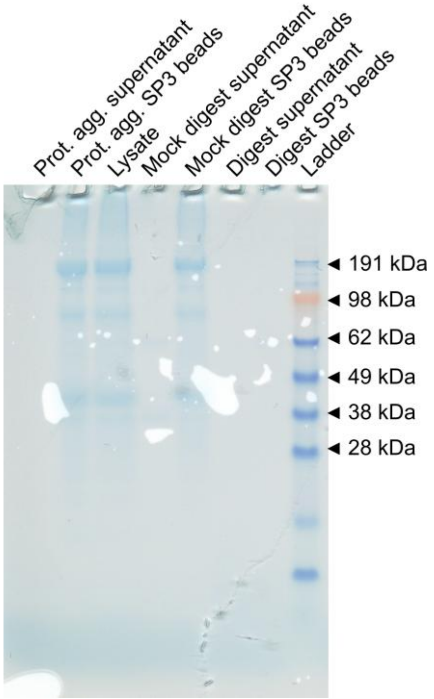


**Figure S3: SP3 digest quality-control gel.** SDS-PAGE of fractions from the SP3 (single-pot, solid-phase-enhanced sample preparation) protein clean-up and digestion workflow, run on mouse macrophage lysate left over from another project rather than on the samples in this study, to check that proteins are captured on the beads and peptides are released. Lanes, left to right: protein-aggregation supernatant, protein-aggregation SP3 beads, input lysate, mock-digest supernatant, mock-digest SP3 beads, digest supernatant, digest SP3 beads, and molecular-weight ladder.

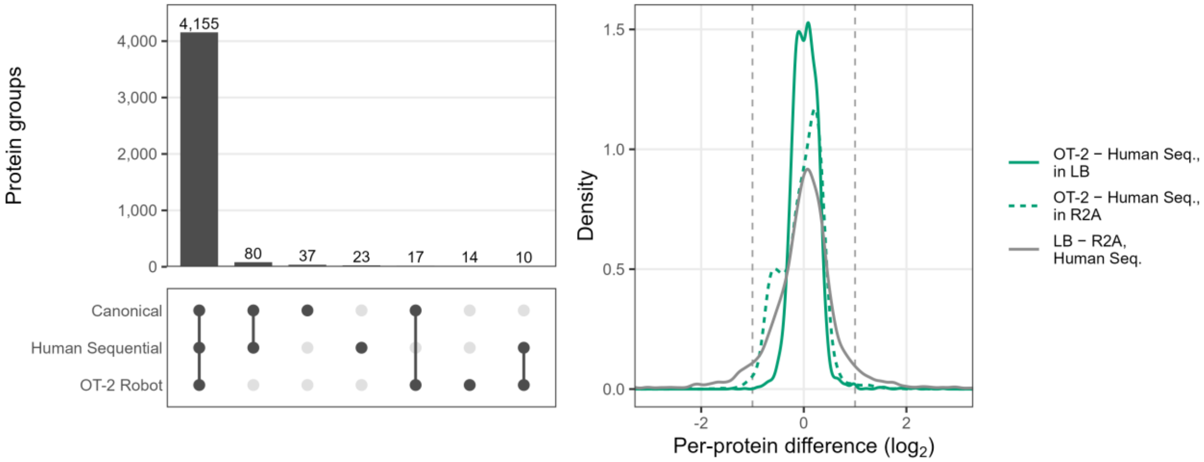


**Figure S4: The three methods quantify the same proteins, with no evidence of selective loss.** (A) Overlap between methods of protein groups quantified in all four replicates of at least one medium; 4,155 of 4,336 are shared by all three methods. All 80 groups complete in both manual methods but not in the OT-2 Robot set were still detected in OT-2 Robot samples, and 31 of them were missing only from the flagged replicate (Figure S5). (B) Per-protein difference in group mean (median-centered $\log_2$ MaxLFQ intensity) for the 3,923 protein groups quantified in every sample. OT-2 Robot minus Human Sequential centres on zero in LB and R2A (medians 0.03 and 0.02), with 0.5% and 2.1% of protein groups beyond $\pm 1$ $\log_2$ (dashed lines), against 11.5% for LB minus R2A within Human Sequential. The R2A trace includes the flagged replicate.

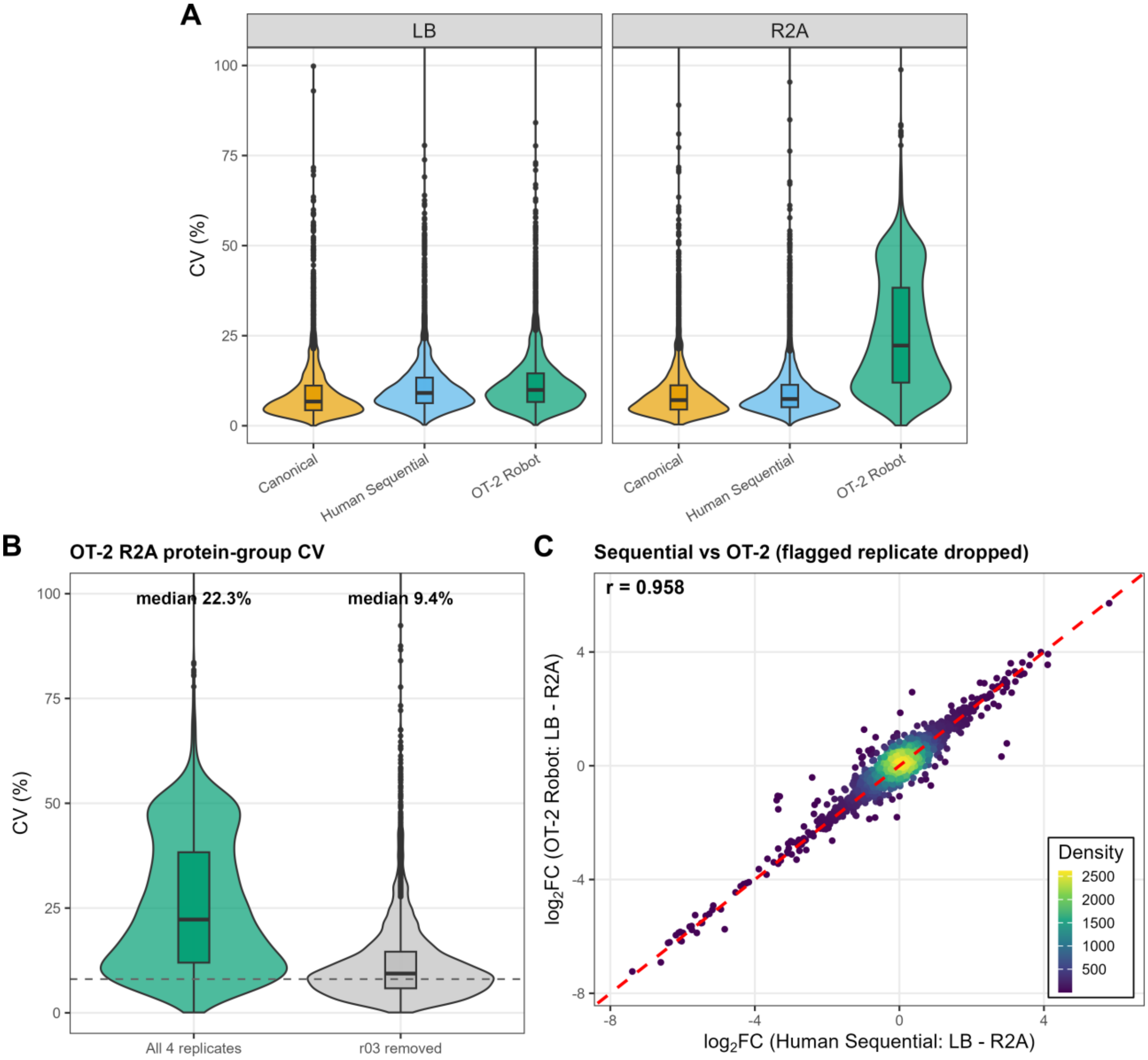


**Figure S5: Proteomics replicate precision and outlier sensitivity.** (A) Coefficient of variation (CV%) of QuantUMS/MaxLFQ protein-group intensities across the four biological replicates within each method × media group; CVs are comparable except for OT-2 R2A. (B) OT-2 R2A protein-group CV with all four replicates (median 22.3%) versus the flagged replicate removed (median 9.4%, in line with the other five method × media groups; dashed line). The flagged replicate carries ~28% less total signal and correlates 0.97 with its groupmates against 0.997 among the other three, consistent with an Evotip loading error downstream of the extraction; the lipid and metabolite fractions from the same extraction are unremarkable. (C) Per-protein $\log_2$ fold change (LB − R2A), Human Sequential versus OT-2 Robot, recomputed with the flagged replicate dropped: the Pearson correlation ($r = 0.958$) is essentially unchanged from the full-data value ($r = 0.939$; main-text Figure 2 D). The replicate is retained in all analyses; removing it changes none of the conclusions.

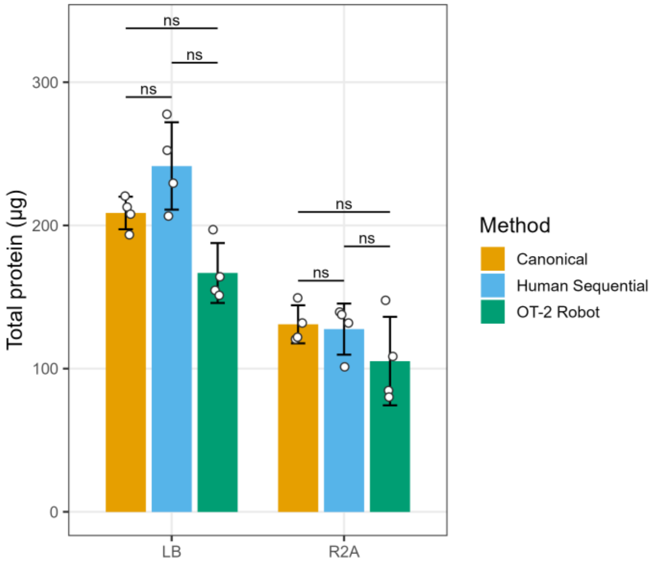


**Figure S6: Total protein recovered by extraction method.** Pierce BCA quantification of the protein fraction recovered from each *P. putida* extraction (n = 4 per method × media; bars, group mean; error bars, SD; points, individual replicates; brackets, Welch's *t*-tests between methods within each medium, Holm-adjusted across the six comparisons; ns, adjusted $P \geq 0.05$). The OT-2 Robot recovered 69–83% of the manual yields on average, and no pairwise difference within a medium was significant after correction (smallest adjusted $P = 0.052$, OT-2 Robot versus Human Sequential in LB). LB exceeded R2A for every method (Welch's *t*-test, $P \leq 0.02$).

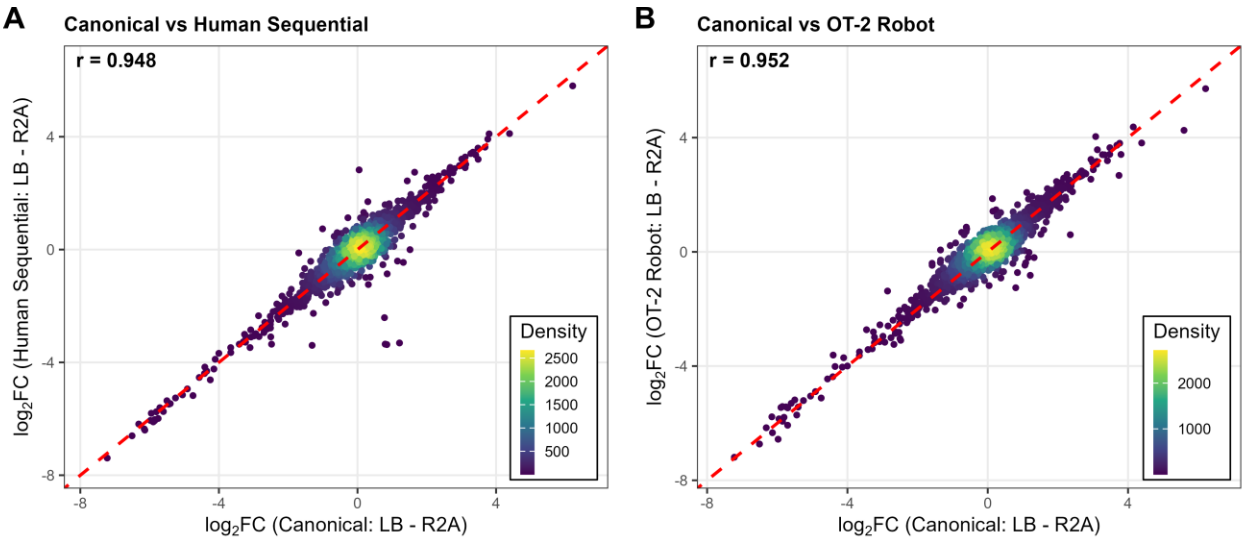


**Figure S7: Proteomics fold-change concordance for the other method comparisons.** Per-protein $\log_2$ fold change (LB − R2A) compared between (A) Canonical and Human Sequential, isolating the chemistry change (r = 0.948), and (B) Canonical and OT-2 Robot, the full modified-and-automated workflow (r = 0.952). Point density is color-coded; the red dashed line is the identity. The Human Sequential versus OT-2 Robot comparison is shown in main-text Figure 2 D.

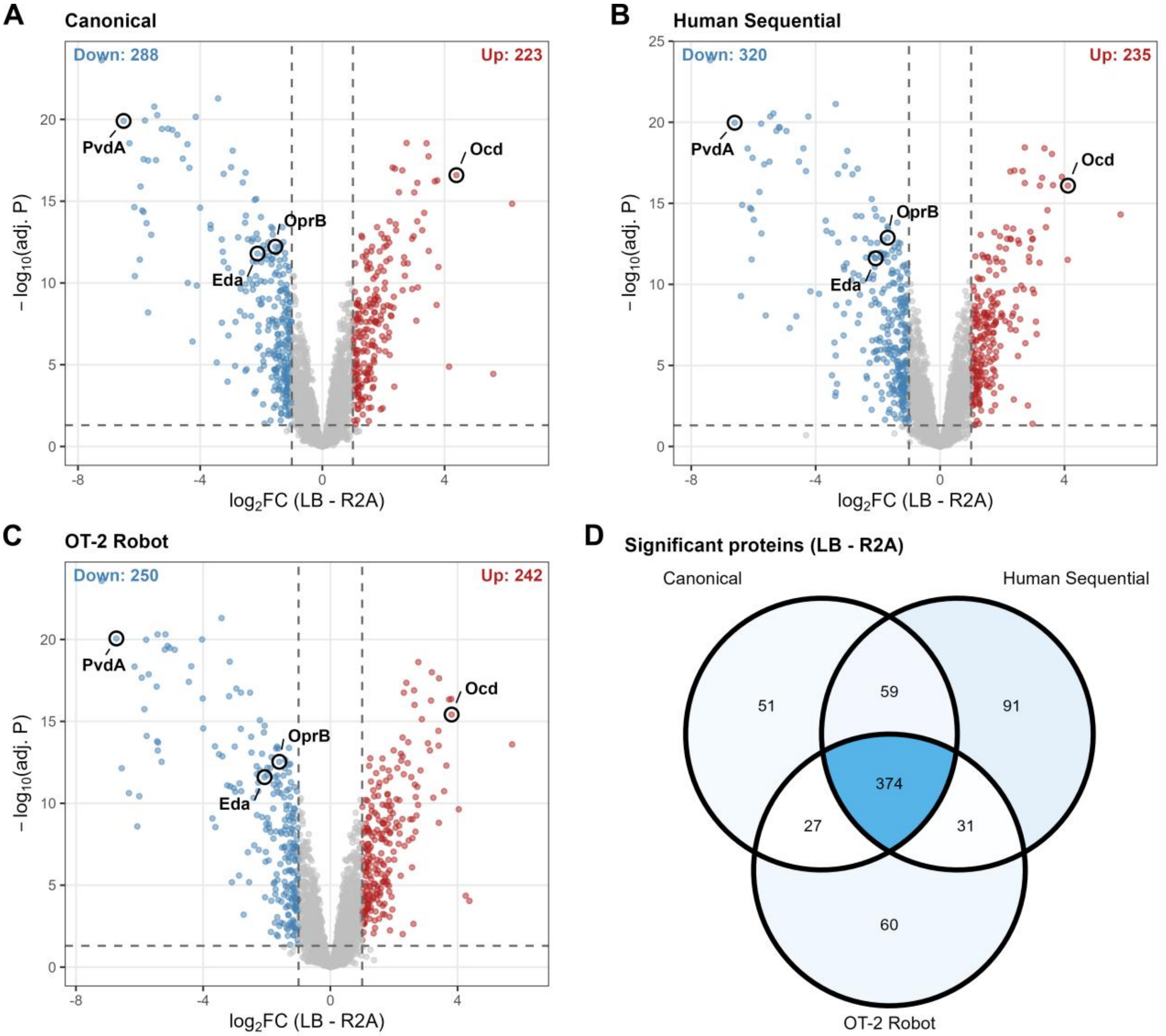


**Figure S8: Differential abundance between media is consistent across methods.** (A–C) Volcano plots of per-protein $\log_2$ fold change versus $-\log_{10}$ adjusted $P$ (limma moderated $t$-test, LB vs R2A) for Canonical, Human Sequential, and OT-2 Robot extractions; dashed lines mark adjusted $P = 0.05$ and $|\log_2 \mathrm{FC}| = 1$, with counts of significantly up- and down-regulated proteins annotated. Four marker proteins are labeled in every panel: OprB (glucose-uptake porin) and Eda (Entner–Doudoroff KDPG aldolase), higher in R2A; PvdA (pyoverdine biosynthesis, the iron-starvation response and the single largest media signal), also higher in R2A; and Ocd (ornithine cyclodeaminase, arginine catabolism), higher in LB. All four shift in the same direction in every method. (D) Venn diagram of the protein groups called significant (adjusted $P < 0.05$, $|\log_2 \mathrm{FC}| > 1$) in each method; the shared core of 374 protein groups shows the three methods recover the same differentially abundant proteins.

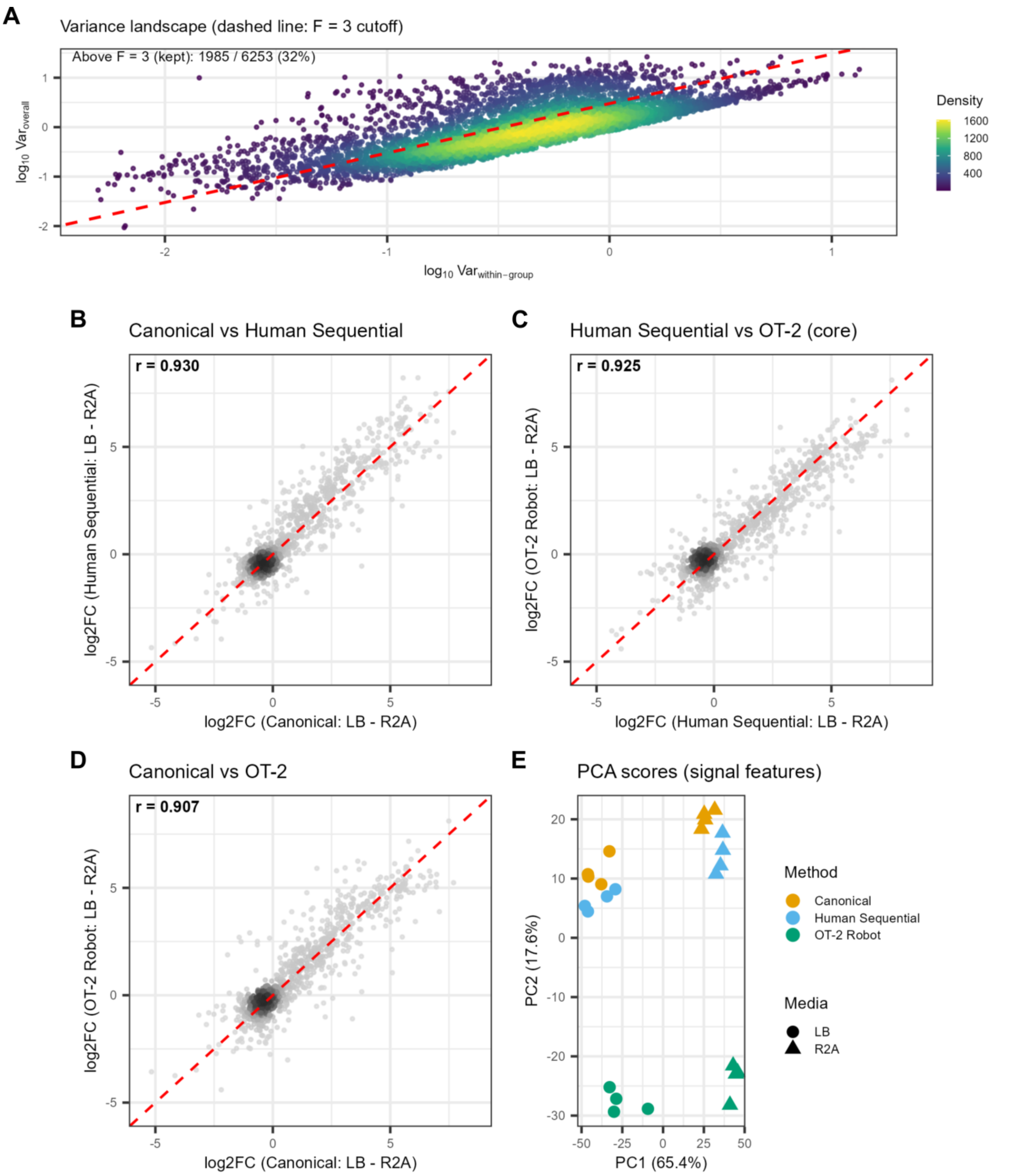


**Figure S9: Metabolomics method-equivalence sanity check.** (A) Per-feature variance landscape with the $F_{\text{within}} = 3$ cutoff; 1,985 / 6,253 features (32%) are retained as signal features. (B–D) Per-feature $\log_2$ fold change (LB − R2A) compared across methods: Canonical versus Human Sequential (r = 0.930), Human Sequential versus OT-2 Robot (r = 0.925), and Canonical versus OT-2 Robot (r = 0.907). (E) PCA scores of signal features; PC1 separates LB from R2A and PC2 separates OT-2 Robot from the manual methods. Concordance is uniformly high (r = 0.907–0.930), consistent with the three methods sharing identical aqueous-phase chemistry. No internal standards or pooled QC samples were used for the metabolomics fraction, and the four blanks were not used in the analysis; this dataset is presented as a method-equivalence check rather than a discovery analysis.

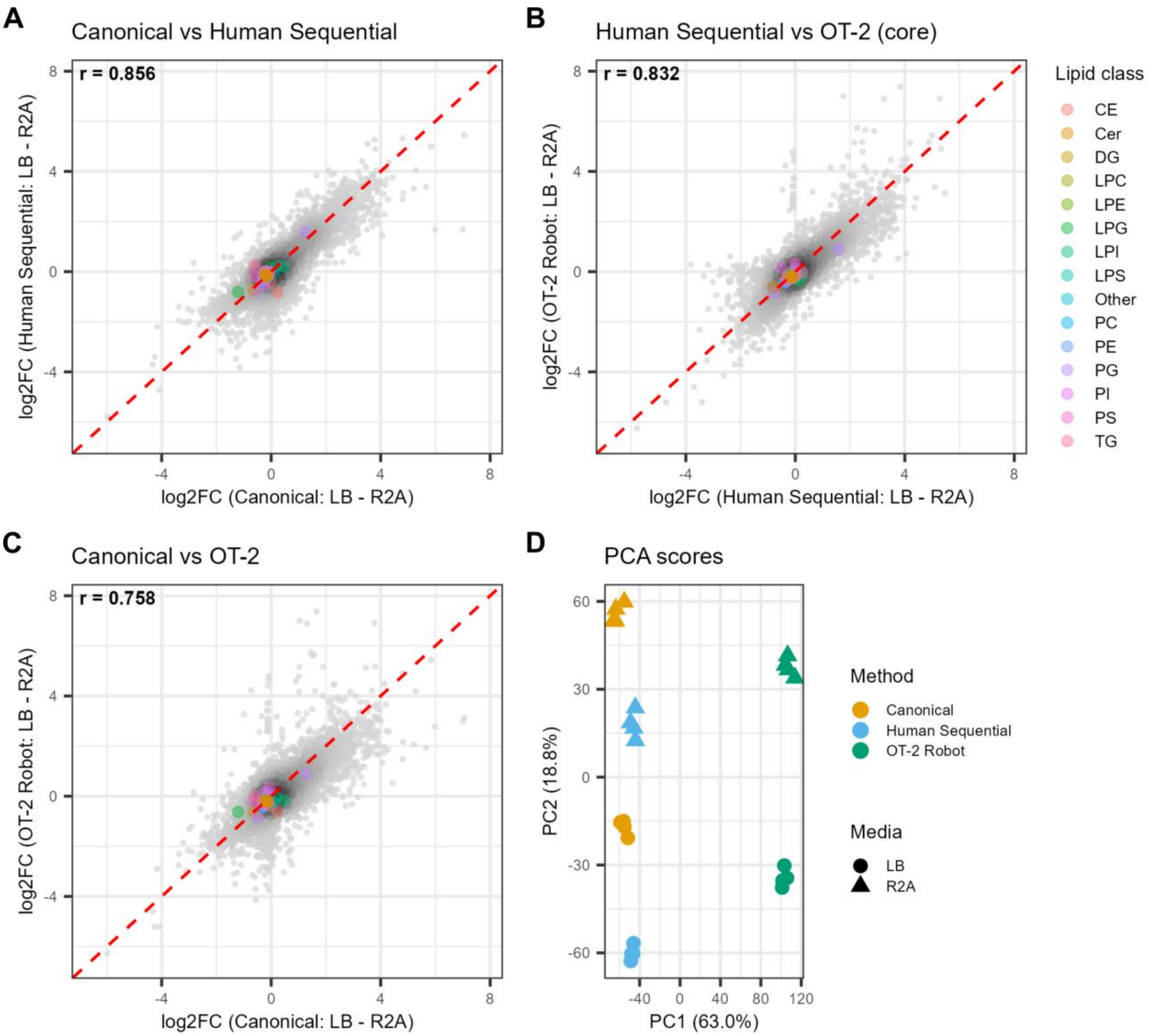


**Figure S10: Lipidomics method equivalence.** (A–C) Per-feature $\log_2$ fold change (LB − R2A) compared across methods: Canonical versus Human Sequential (r = 0.856), Human Sequential versus OT-2 Robot, isolating automation (r = 0.832), and Canonical versus OT-2 Robot (r = 0.758). Background grayscale density, signal features ($F_{\text{within}} > 3$); colored points, SPLASH internal standards by lipid class (constant-concentration spike-ins that cluster at the origin and serve as analytical anchors only); Pearson r is computed on the signal-feature background. (D) PCA scores of signal features. PC1 (63%) separates samples by extraction method. Because the lipidomics samples were acquired in method-blocked run order, extraction method and run order are confounded by design, so PC1 reflects this combined axis. PC2 (18.8%) separates LB from R2A within every method cluster.

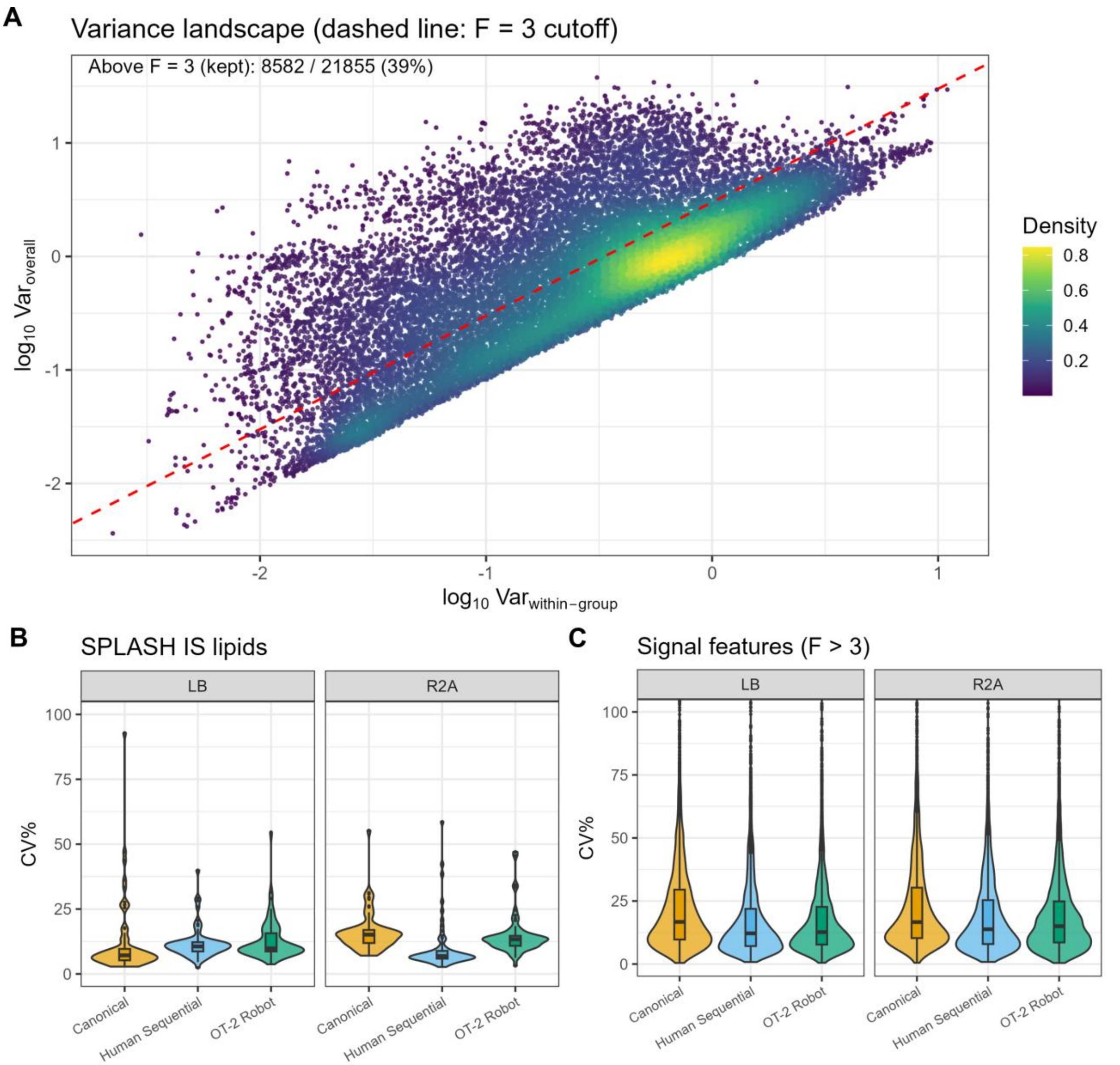


**Figure S11: Lipidomics reproducibility.** (A) Per-feature variance landscape: $\log_{10}$ within-group variance versus $\log_{10}$ overall variance, density-colored. The dashed red line marks the $F_{\text{within}} = 3$ cutoff; 8,582 / 21,855 features (39%) sit above the line and were retained as signal features. (B) CV% of the 69 SPLASH internal-standard lipids, faceted by media, which gives the analytical floor (median ≈ 10%). (C) CV% of complete-case signal features ($F_{\text{within}} > 3$), faceted by media. The additional variance over the analytical floor is consistent across all three methods, indicating equivalent extraction precision.

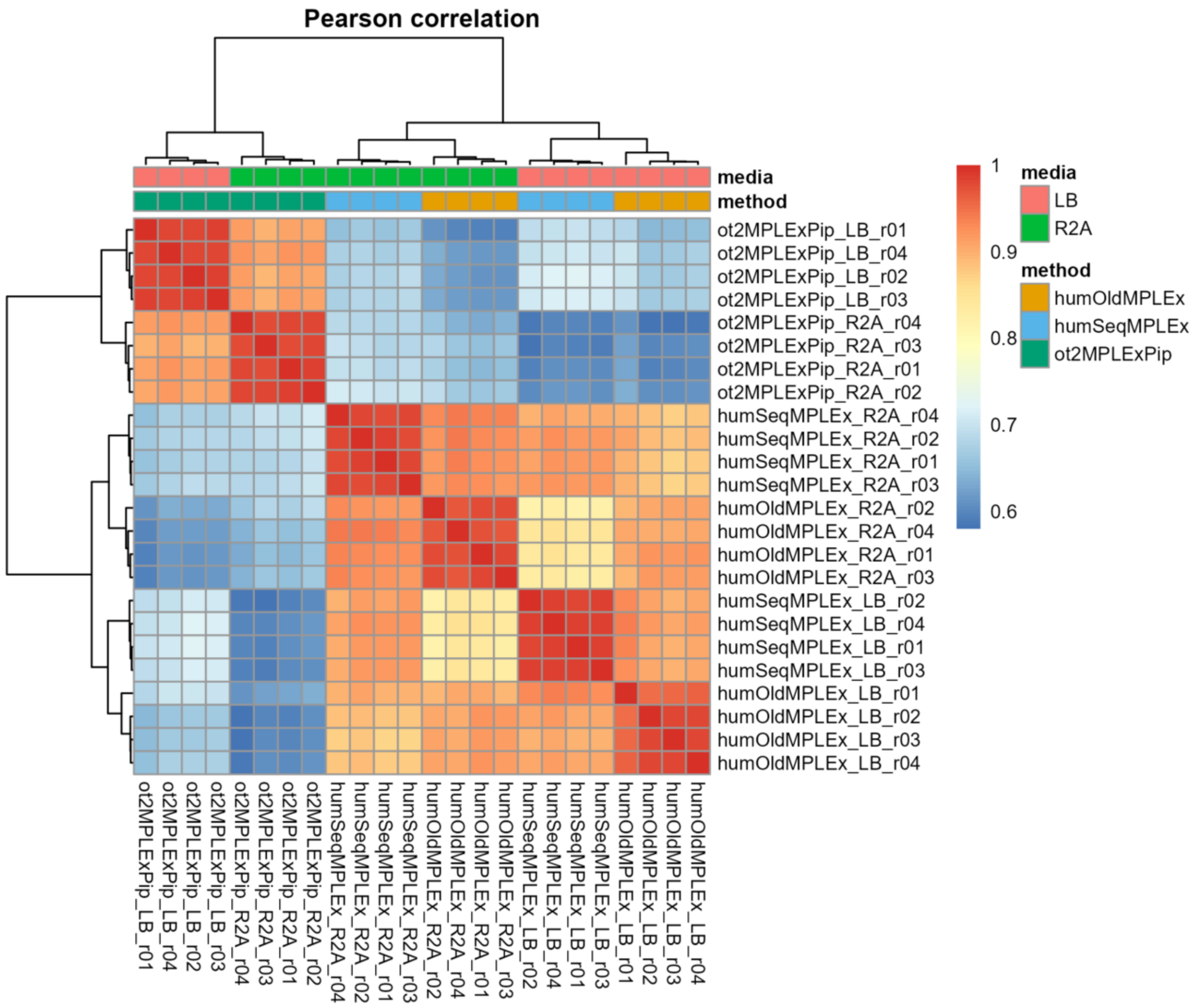


**Figure S12: Lipidomics sample correlation.** Pairwise Pearson correlation of signal-feature intensities ($F_{\text{within}} > 3$) across all 24 lipidomics samples, with hierarchical clustering on both axes. Annotation bars indicate extraction method and growth media. Sample names use internal method labels: humOldMPLEx, Canonical; humSeqMPLEx, Human Sequential; ot2MPLExPip, OT-2 Robot. Samples separate first into OT-2 Robot and the two manual methods, and then by media within each branch. Extraction method and run order are confounded in these samples, so the first split reflects both (Figure S10).

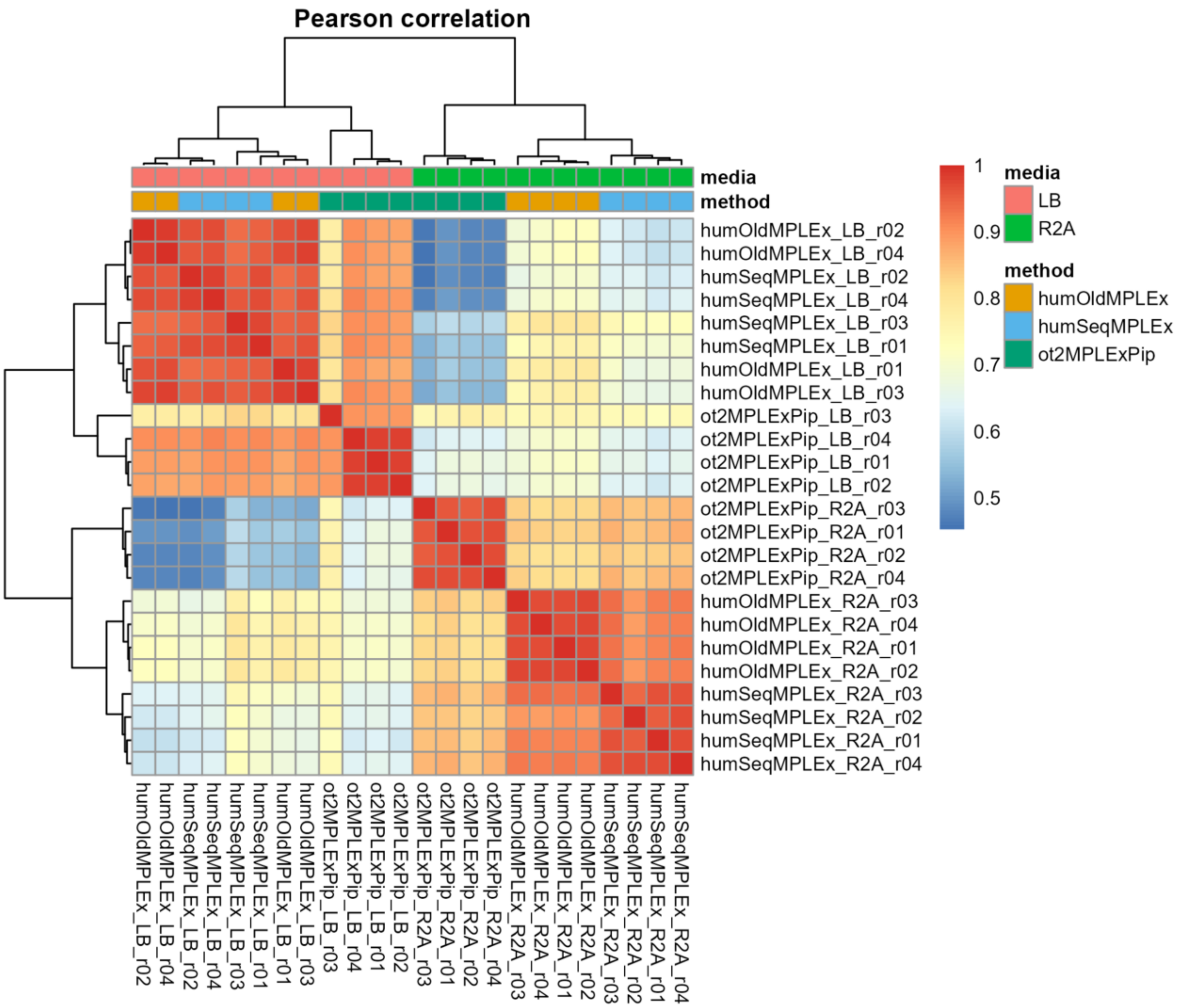


**Figure S13: Metabolomics sample correlation.** Pairwise Pearson correlation of signal-feature intensities ($F_{within} > 3$) across all 24 metabolomics samples, with hierarchical clustering on both axes. Annotation bars indicate extraction method and growth media.

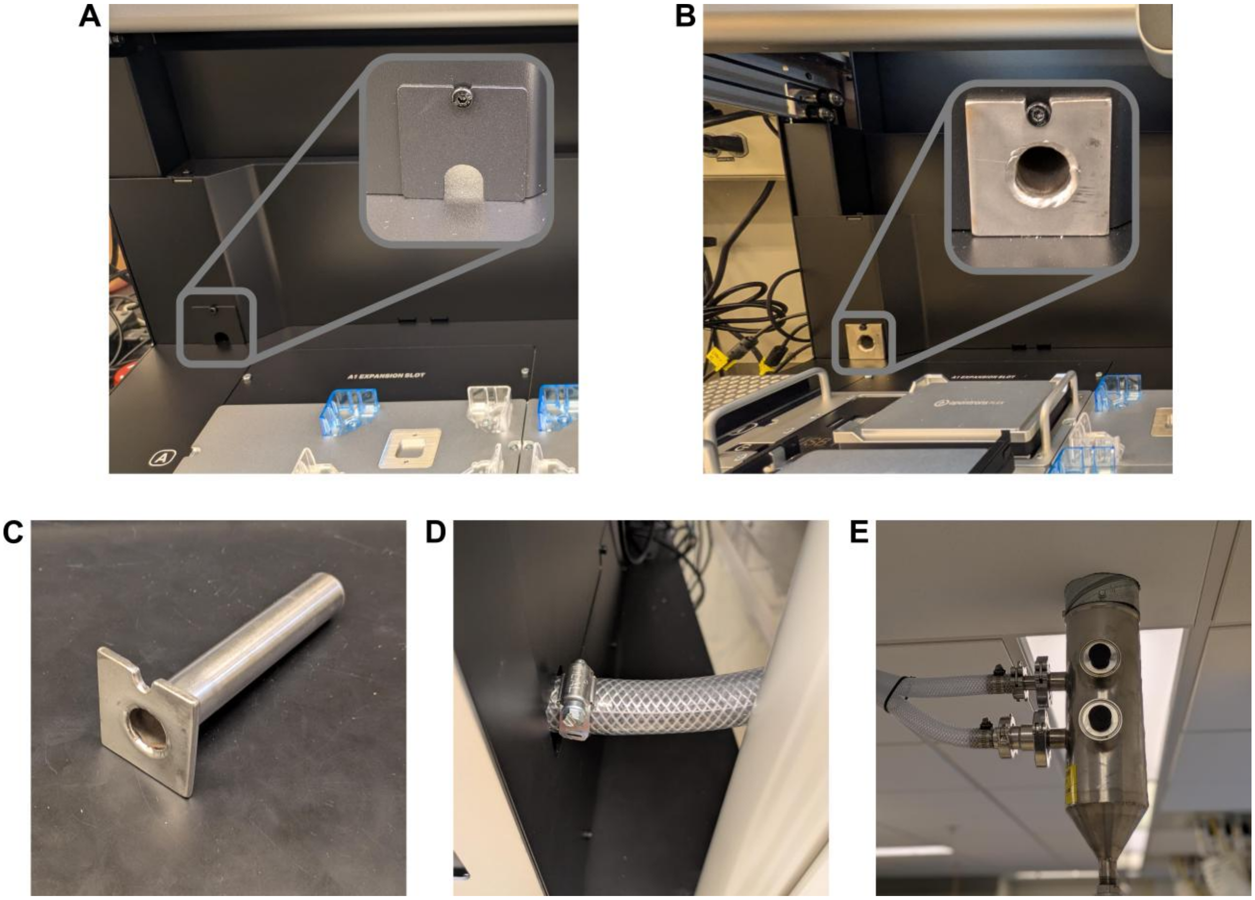


**Figure S14: Ventilation of the Opentrons Flex.** (A) Manufacturer's cover plate over the cable pass-through port on the rear wall of the Flex deck (inset: magnified view). (B) The same port with the cover plate replaced by a custom ventilation adaptor (inset: magnified view). (C) Custom-built stainless steel ventilation conduit, consisting of a mounting plate welded to a tube. The conduit was fabricated by our in-house machine shop by tracing the manufacturer's cover plate onto stainless steel stock, cutting it to shape, and welding on the tube; no CAD files or technical drawings exist. (D) The ventilation conduit passes entirely through the rear panel of the Flex and connects to reinforced flexible tubing secured with a hose clamp. (E) The tubing connects to a ceiling-mounted stainless steel house-vacuum line via sanitary clamp fittings.

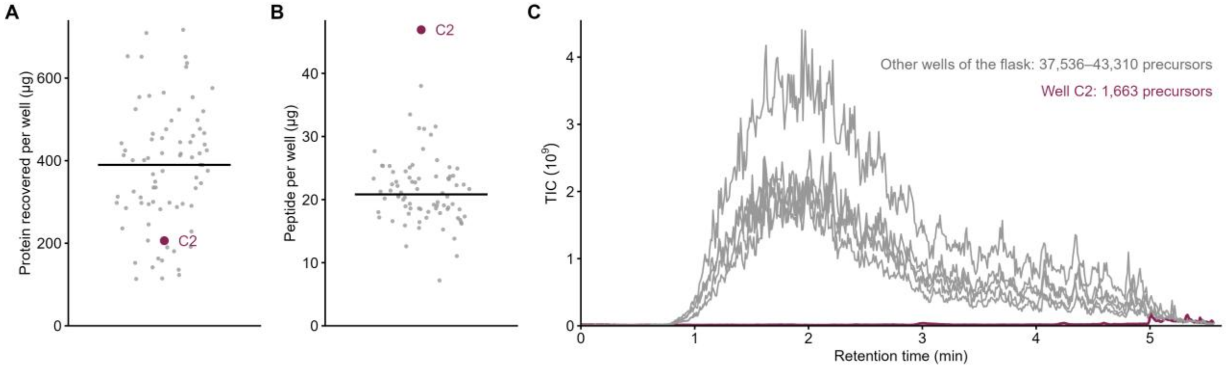


**Figure S15: Well C2 carried sample through digestion and was lost to a sample handling error before LC-MS.** (A) Protein recovered per well after extraction and (B) peptide per well after SP3, both by BCA, for all 80 wells (line, plate median; wine, well C2). C2 recovered 206 µg of protein, was normalized to 50 µg for SP3 like the other wells, and read 47 µg of peptide. (C) TIC of C2 against the other five wells from the same flask. C2 is blank (1,663 precursors, against 37,536–43,310) because of a sample handling error after the peptide BCA and upstream of the LC-MS, rather than a fault in extraction or SP3.

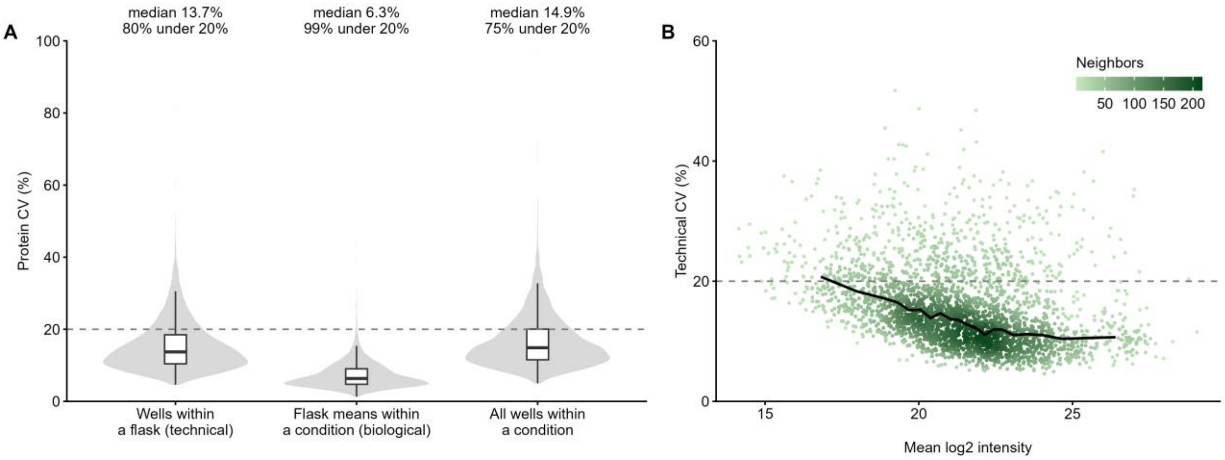


**Figure S16: Full-workflow precision on the carbon × nitrogen plate, at the technical and biological levels.** (A) Per-protein coefficient of variation (CV, linear scale) for the 2,881 protein groups quantified in all 79 wells, at three levels: across the 5–7 wells of one flask (technical; each well is an independent run of the full workflow, from extraction to LC-MS; median of the 12 flasks), across the three flask means of a condition (biological; median of the four conditions), and across all wells of a condition ignoring flask (the replicate CV most studies report; median of the four conditions). Labels give the median CV (13.7%, 6.3% and 14.9%) and the share of protein groups under 20% (dashed line). Each flask mean averages 5–7 wells, so the biological CV carries only a fraction of the technical noise and mostly reflects culture-to-culture variation. (B) Technical CV against mean $\log_2$ intensity, colored by the number of neighboring proteins; the line is the median in 20 abundance bins. Two protein groups with technical CV > 60% lie outside the plotted range. Per-condition technical CVs are shown in main-text Figure 4 D.

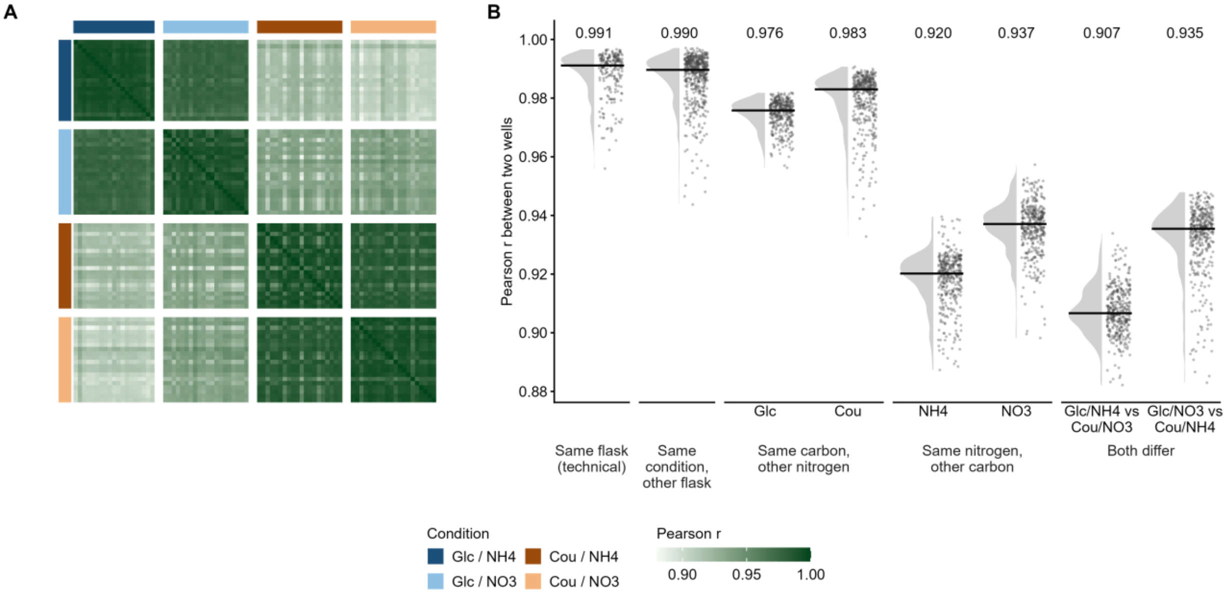


**Figure S17: Sample-to-sample correlation on the carbon × nitrogen plate.** (A) Pearson *r* between all pairs of the 79 wells, computed on the 2,881 protein groups quantified in every well (median-centered $\log_2$ MaxLFQ intensities). Wells are in design order (condition, then flask), and bars mark condition. (B) The same 3,081 pairs grouped by how the two wells are related, with the median *r* above each group: two preps from the same flask (technical replicates, 0.991), wells from different flasks of the same condition (0.990), and pairs that differ in one or both nutrients. The classes that cross a nutrient are split by the conditions involved, because their pairs are not equivalent. Nitrogen changes the glucose proteome more than the coumarate proteome (0.976 vs 0.983), and carbon changes the proteome less under nitrate than under $NH_4$ (0.937 vs 0.920). This is the carbon × nitrogen interaction. Half-violins show each group's distribution, and points are individual pairs. Pairs share wells, so the distributions are descriptive rather than independent tests.

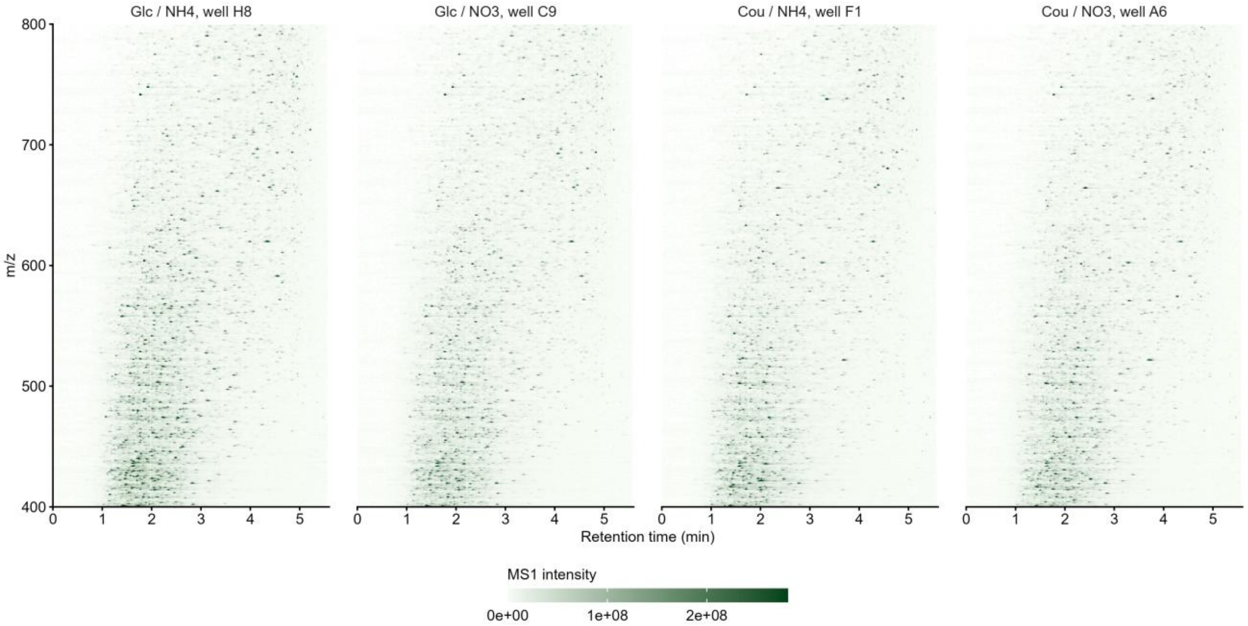


**Figure S18: MS1 maps show no polymer contamination in the proteomics samples.** Absolute MS1 intensity by retention time and $m/z$ for the median-signal well of each condition, summed in 0.02 min × 0.5 Th bins. The color scale is linear and tops out at the 99.9th percentile of bins. Polymer contaminants would appear as a repeating ion series rising in $m/z$ with retention time; none is visible.

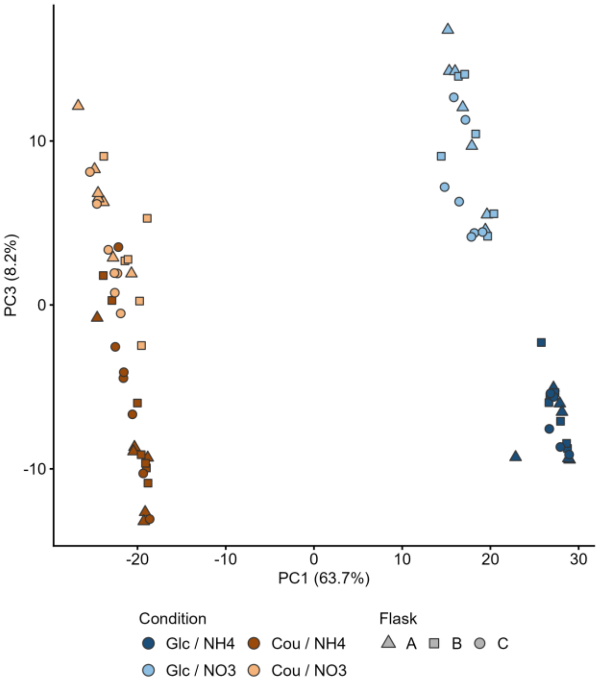


**Figure S19: PCA of the carbon × nitrogen plate, PC1 against PC3.** The same principal component analysis as main-text Figure 4 E (2,881 protein groups quantified in all 79 wells; median-centered $log_2$ MaxLFQ intensities), with PC3 (8.2% of the variance) in place of PC2 (12.0%). Point = well (technical replicate); fill = condition; shape = flask (biological replicate). PC1 (63.7%) separates the carbon sources. Within each carbon source, PC3 separates $NH_4$ from $NO_3$ by 5.0 (glucose) and 2.5 (coumarate) pooled within-group standard deviations, against 1.4 and 0.8 on PC2.

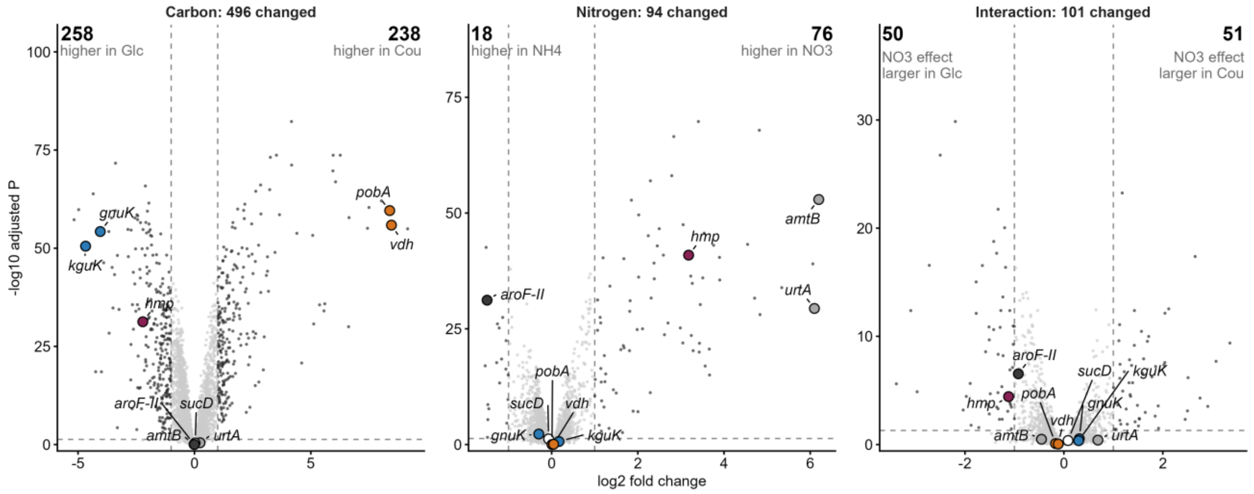


**Figure S20: Differential abundance on the carbon × nitrogen plate.** Volcano plots for the carbon (coumarate − glucose, averaged over nitrogen), nitrogen ($NO_3 - NH_4$, averaged over carbon) and interaction contrasts (limma, with wells from the same flask treated as correlated so that the flask is the replicate). Dark points, called changed (Benjamini–Hochberg adjusted $P < 0.05$ and $|\log_2 FC| > 1$); bold numbers, count on each side; highlighted, the proteins from Figure 4 F, with the same fills. Protein groups absent from a whole condition have no fold change and are not shown.

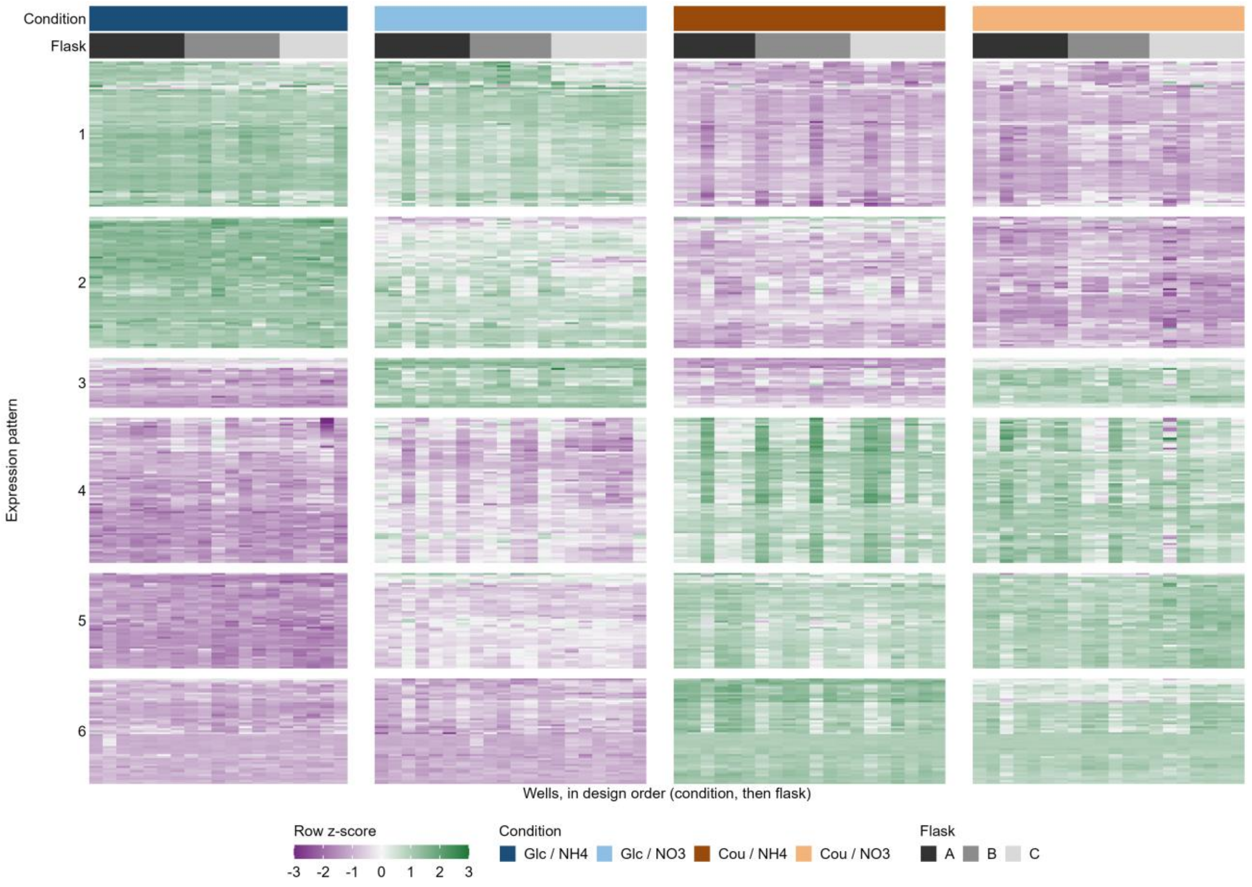


**Figure S21: Proteins that respond to carbon or nitrogen, well by well.** Row z-scores of median-centered $\log_2$ MaxLFQ intensities for the 338 protein groups that are quantified in all 79 wells and called changed in at least one contrast (carbon, nitrogen or carbon × nitrogen; limma with the flask as the replicate; adjusted $P < 0.05$ and $|\log_2 FC| > 1$). Columns are wells, each an independent run of the full workflow, ordered by condition and then flask (bars above). Rows are clustered (Ward linkage, Euclidean distance), and the tree is cut into six expression patterns, numbered from the top. Patterns 1 and 2 are higher in glucose, pattern 2 most strongly with

$NH_4$; patterns 4–6 are higher in coumarate, pattern 6 most strongly with $NH_4$; pattern 3 is higher in nitrate under both carbon sources. z-scores are clipped at ±3. The rows were selected for changing with condition, so the figure shows the shape of the response rather than testing it. Proteins quantified under only one nitrogen source (e.g., NasA, NirD) are absent because they are not quantified in every well.

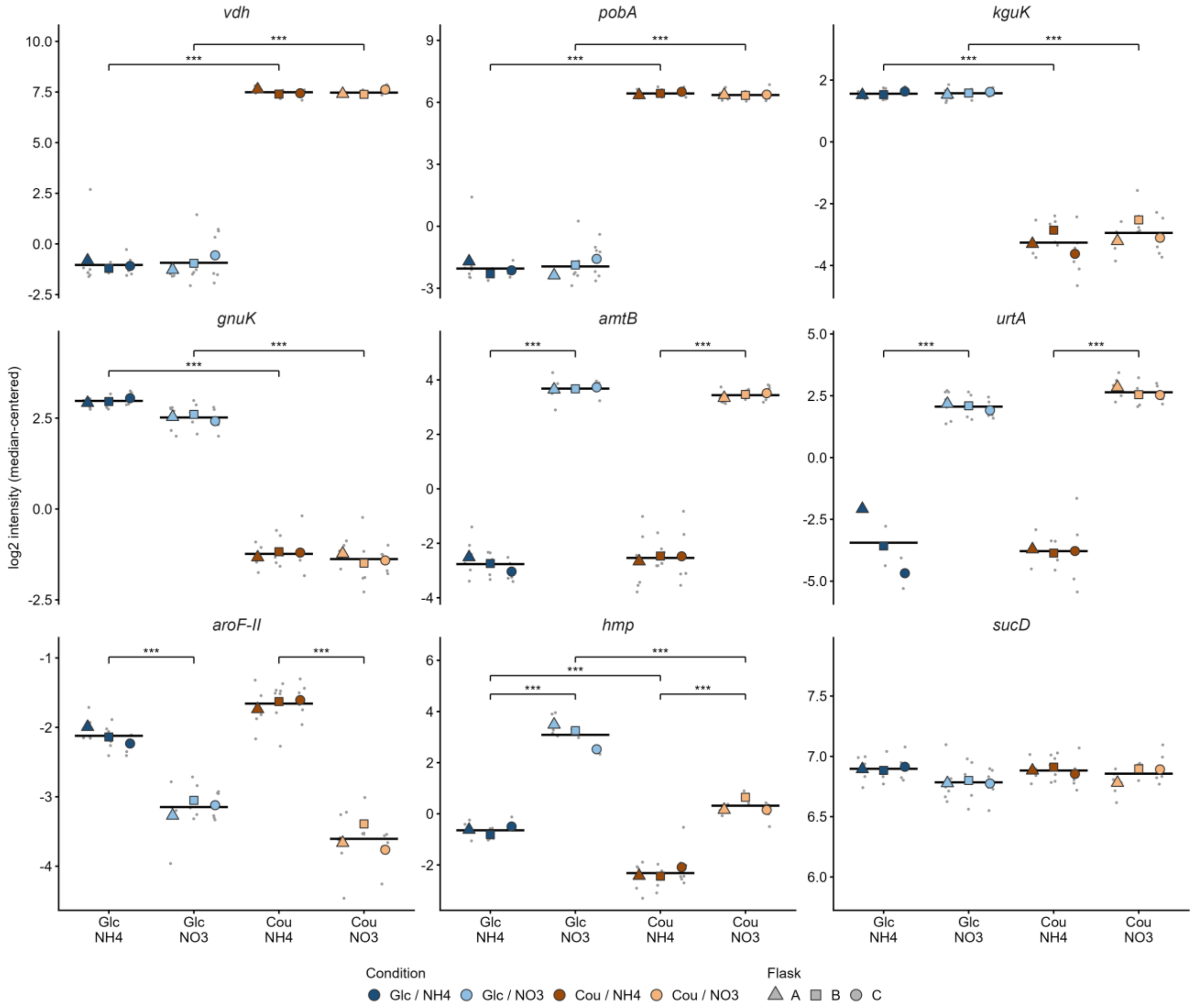


**Figure S22: Representative proteins from main-text Figure 4 F.** The two largest coumarate-responsive (*vdh*, *pobA*), glucose-responsive (*kguK*, *gnuK*) and nitrate-responsive (*amtB*, *urtA*) proteins, the largest $NH_4$-responsive protein (*aroF-II*), the protein most changed by both nutrients (*hmp*), and an unchanged reference (*sucD*). Small gray points, wells (technical replicates); large symbols, flask means (biological replicates; fill = condition, shape = flask); black bars, condition means (mean of the three flask means). Values are median-centered $\log_2$ MaxLFQ intensities; y-axes differ between panels, and each spans at least 2 $\log_2$ units. Brackets mark single-nutrient comparisons (nitrogen within each carbon source, carbon within each nitrogen source) that pass the main analysis's call rule: Benjamini–Hochberg-adjusted $P < 0.05$ and $|\log_2 \text{FC}| > 1$ (limma, with wells from the same flask treated as correlated; * $P < 0.05$, ** $P < 0.01$, *** $P < 0.001$). *sucD* changes by at most 0.11 $\log_2$ in any comparison. Missing points are wells in which the protein was not quantified (*urtA* is quantified in 15 of 39 $NH_4$ wells).

#### Supplemental Tables

**Table S1:** Per-method lipidomics summary statistics. Signal features pass $F_{within} > 3$ across the entire dataset and are the same for all rows, because they come from a single feature pool. Median CV is reported separately for SPLASH internal standards and endogenous signal features. Differential features are those differentially abundant between LB and R2A (Welch's *t*-test, BH-adjusted $p < 0.05$, $|\log_2 FC| > 1$).

| Method | Complete-case features | Signal features | Median CV, SPLASH (%) | Median CV, endogenous (%) | Differential features |
|---|---|---|---|---|---|
| Canonical | 5772 | 8582 | 11.6 | 16.7 | 1392 |
| Human Sequential | 5772 | 8582 | 8.8 | 12.9 | 1509 |
| OT-2 Robot | 5772 | 8582 | 11.8 | 13.7 | 1364 |
